# Maternal sucrose intake during pregnancy alone disrupts testosterone and allopregnanolone levels in the fetal brain of rats

**DOI:** 10.1101/2025.07.21.664469

**Authors:** Minseon M. Jung, Marwa Idrissi, Yen-Nhi Hoang, Tao Huan, Désirée R. Seib, Kiran K. Soma

## Abstract

**Background:** Intake of added sugars, such as sucrose, is high globally. In rats, a maternal high-sucrose diet (HSD) from 10 wk before pregnancy to embryonic day (E)19.5 has widespread impacts on maternal, placental, and fetal blood and brain steroid levels, including glucocorticoids, androgens, and aldosterone.

**Objective:** This study examined whether maternal HSD during pregnancy alone is sufficient to alter maternal, placental, and fetal steroids.

**Methods:** Pregnant rats received either a control diet (1% kcal sucrose) or an isocaloric, nutrient-matched HSD (26% kcal sucrose) between E0.5-19.5. On E19.5, we collected maternal serum, placenta, fetal blood and brain, and amniotic fluid. We microdissected the placenta and fetal brain and measured 14 steroids using highly sensitive and specific liquid chromatography-tandem mass spectrometry (n=12-15/diet/sex).

**Results:** Maternal HSD during pregnancy alone did not alter maternal food intake, maternal body mass, and litter size (all p values ≥ 0.29, Student’s t-test) but increased the percentage of males in a litter (p = 0.03, Student’s t-test). Maternal HSD did not alter steroids in the maternal serum (all p values ≥ 0.21, Student’s t-test), placenta (all p values ≥ 0.07, 2-way analysis of variance (ANOVA)), and fetal blood (all p values ≥ 0.13, 2-way ANOVA). Nonetheless, maternal HSD increased testosterone in the fetal nucleus accumbens (p = 0.04, 2-way ANOVA), decreased allopregnanolone in the fetal amygdala (p = 0.01, 2-way ANOVA), and decreased 11-dehydrocorticosterone in the amniotic fluid (p = 0.05, 2-way ANOVA).

**Conclusions:** Maternal HSD during pregnancy alone does not affect steroid levels in the maternal serum, placenta, or fetal blood, but disrupts testosterone and allopregnanolone levels in critical regions of the fetal brain that regulate reward-seeking and emotion. Thus, while a long-term maternal HSD is necessary for widespread endocrine effects, the fetal brain is sensitive to short-term increases in maternal sucrose consumption during pregnancy.

## INTRODUCTION

Added sugar intake around the world exceeds recommendations (<10% of total energy intake) (1,2). In particular, women increase sweetened food consumption during pregnancy (3). High maternal sugar intake is associated with pregnancy complications and gestational weight gain (4), as well as increased body mass index and weight in children (5,6). Conversely, exposure to sugar rationing *in utero* and during early life reduces the risk of Type II diabetes and hypertension (7). Thus, maternal sugar intake impacts offspring health.

Sucrose (table sugar) is a common added sugar in processed foods and beverages. The metabolic and cardiovascular effects of sucrose intake are well-studied (8,9), but little is known about how sucrose impacts the brain and behaviour. In humans, high sugar intake is associated with depression, addiction-related behaviours, and impulsivity (10,11). In adult rodents, sucrose intake alters synaptic transmission, memory, and locomotion (12–15).

Very few studies have examined how maternal sucrose consumption affects the developing offspring brain, which is more vulnerable than the adult brain. In mice, sucrose exposure *in utero* increases locomotion in adolescent male offspring (16). Perinatal sucrose exposure reduces spatial memory in adolescent and adult female rat offspring (17). Prenatal exposure to sucrose impairs spatial memory (18,19) and increases hippocampal apoptosis (18) in male rat offspring.

There are limitations of previous animal studies investigating the effects of sugar intake on the brain. First, some studies used a standard rodent chow as a control diet to compare to a high-sugar diet, but these two diets also differed in caloric density and macro- and micronutrient composition (20–22). Moreover, some standard rodent chows, such as AIN-76, contain high levels of sucrose (23–25) and are not appropriate control diets. Second, in some studies, sugar was administered in drinking water (12,16,24–26), which reduces food consumption (25–27) and thus decreases intakes of protein, fat, and micronutrients. Third, some studies gave diets with ∼60% kcal from sugar (17,20,22,28), which is far above average human intakes. Lastly, some studies used diets that are high in sugar and fat (21,29), which cannot isolate the effects of sugar.

To address these limitations of previous studies, we created a control diet (CON) and high-sucrose diet (HSD) that are isocaloric and macro- and micro-nutrient matched (**Table 1**) (30–32). In the HSD, 25% of kcal from starch is replaced with sucrose. This HSD contains a human-relevant level of sucrose (33). Moreover, this HSD does not alter food intake or body mass in rats (30,31). Thus, we can assess the effects of maternal HSD, without the confounds of maternal obesity or maternal excess caloric intake.

**Table 1:** Composition of the diets.

| CON |  |  |  | HSD |  |  |
| --- | --- | --- | --- | --- | --- | --- |
| Macronutrient | g | kcal (%) |  | g (%) | kcal (%) |  |
| Protein | 19 | 20 |  | 19 | 20 |  |
| Carbohydrate | 67 | 70 |  | 67 | 70 |  |
| Fat | 4 | 10 |  | 4 | 10 |  |
| Total | 90 | 100 |  | 90 | 100 |  |
| kcal/g | 3.8 |  |  | 3.8 |  |  |
| Ingredient | g | kcal | g / kg diet | g | kcal | g / kg diet |
| Casein | 200 | 800 | 189.6 | 200 | 800 | 189.6 |
| L-Cystine | 3 | 12 | 2.8 | 3 | 12 | 2.8 |
| Corn Starch<sup>1</sup> | 550 | 2200 | 521.3 | 300 | 1200 | 284.4 |
| Maltodextrin 10 | 150 | 600 | 142.2 | 150 | 600 | 142.2 |
| Sucrose<sup>2</sup> | 0 | 0 | 0.0 | 250 | 1000 | 237.0 |
| Cellulose, BW200 | 50 | 0 | 47.4 | 50 | 0 | 47.4 |
| Soybean Oil | 25 | 225 | 23.7 | 25 | 225 | 23.7 |
| Lard | 20 | 180 | 19.0 | 20 | 180 | 19.0 |
| Mineral Mix S10026 | 10 | 0 | 9.5 | 10 | 0 | 9.5 |
| Dicalcium Phosphate | 13 | 0 | 12.3 | 13 | 0 | 12.3 |
| Calcium Carbonate | 5.5 | 0 | 5.2 | 5.5 | 0 | 5.2 |
| Potassium Citrate | 16.5 | 0 | 15.6 | 16.5 | 0 | 15.6 |
| Vitamin Mix V10001<sup>3</sup> | 10 | 40 | 9.5 | 10 | 40 | 9.5 |
| Choline Bitartrate | 2 | 0 | 1.9 | 2 | 0 | 1.9 |
| FD&C Red Dye #40 | 0.025 | 0 | 0.0024 | 0 | 0 | 0.0 |
| FD&C Blue Dye #1 | 0.025 | 0 | 0.0024 | 0.05 | 0 | 0.047 |
| Total | 1055 | 4057 |  | 1055 | 4057 |  |
| Sucrose (g%) | 0.9 |  |  | 25 |  |  |
| Sucrose (kcal%) | 1.0 |  |  | 26 |  |  |

Abbreviations: CON: control diet; HSD: high-sucrose diet.
<sup>1,2</sup> CON and HSD differ in corn starch and sucrose content. In HSD, 25% of kcal from starch is replaced with sucrose.
<sup>3</sup>Vitamin Mix V10001 contains 9.3 g/kg diet of sucrose.

Using this diet paradigm, we previously showed that long-term (∼13 or 16 wk) maternal HSD alters neuroendocrine and behavioural outcomes in the offspring. Maternal HSD starting 10 wk before and during pregnancy has widespread impacts on maternal, placental, and fetal steroid levels at embryonic day (E)19.5 (30). Maternal HSD increased glucocorticoids, steroid hormones that regulate metabolism and stress responses, in the maternal serum. Maternal HSD also decreased placenta mass and placental androgens (androstenedione and testosterone). Further, maternal HSD increased aldosterone, a mineralocorticoid and mediator of stress responses (34,35), in the fetal blood. Similarly, in the fetal brain, maternal HSD increased aldosterone and decreased testosterone in the nucleus accumbens (NAc), a critical node in the “reward circuit”. Furthermore, maternal HSD increased the 11-dehydrocorticosteorne (DHC)/corticosterone ratio in several regions of the fetal brain. These data suggest that maternal sucrose consumption is a stressor that disrupts the endocrine system.

Additionally, long-term maternal HSD starting 10 wk pre-pregnancy, during pregnancy, and lactation increases baseline blood and brain corticosterone levels in adult female, but not male, offspring (32). Maternal HSD increases motivation for a sugar reward and increases preference for high-fat diet and high-sucrose diet in adult male offspring (32). Thus, maternal HSD has multiple effects on offspring hormones, brain, and behaviour in adulthood. However, it is unclear whether long-term maternal sucrose consumption is necessary to impact the offspring, or whether maternal sucrose consumption during pregnancy alone is sufficient.

In this study, we used our diet model to determine whether short-term (∼3 wk) maternal HSD during pregnancy alone is sufficient to alter maternal, placental, and fetal steroids. We fed pregnant rat dams either the CON or HSD from E0.5 to E19.5. On E19.5, we collected maternal, placental, and fetal samples. We microdissected the fetal brain, including the NAc and ventral tegmental area (VTA), which are nodes of the mesocorticolimbic “reward” circuit. We also examined the amygdala (AMY) and ventral hippocampus (vHPC), which are implicated in emotion and anxiety. Lastly, we examined the hypothalamus (HYP), which is important for homeostasis and the stress response. We measured 14 steroids, including glucocorticoids, progestins, androgens, estrogens, and aldosterone, using ultrasensitive and specific liquid-chromatography tandem mass spectrometry (LC-MS/MS) (**Figure 1**).

**Figure 1:**
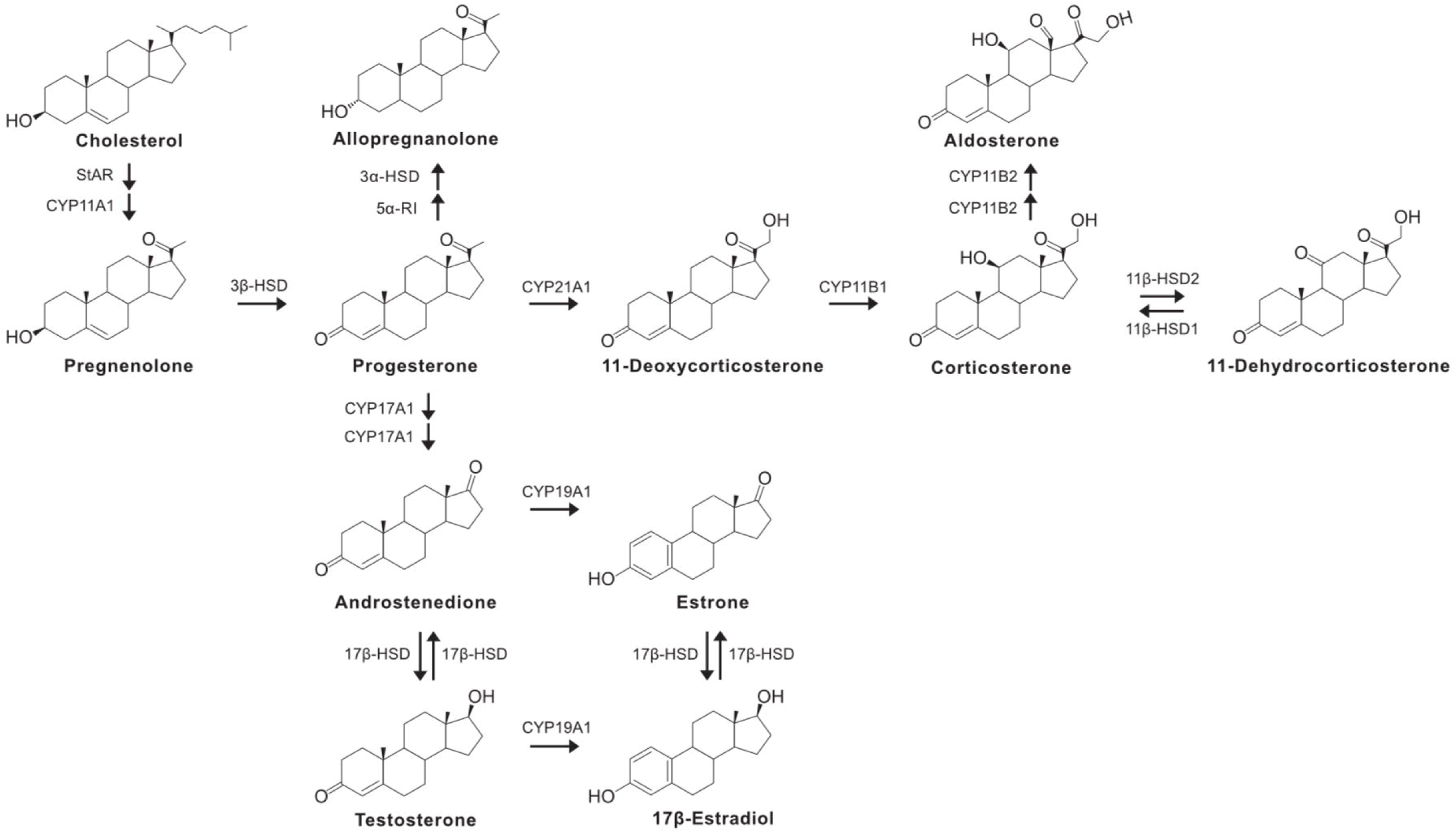
Steroids for LC-MS/MS analysis. A simplified steroidogenic pathway that illustrates the steroids that were examined by LC-MS/MS in this study. Cholesterol was not measured. LC-MS/MS: liquid chromatography-tandem mass spectrometry.

## MATERIALS AND METHODS

### Animals

Adult female Long-Evans rats (n=16/diet; postnatal day 57-63, 188-262 g; Charles River Laboratories, Kingston, NY, USA) were pair housed with *ad libitum* access to standard rodent chow (Rat diet 2918; Harlan Teklad Global, Madison, WI, USA) and water (reverse osmosis purification and chlorination sterilization). All animals were housed in clear, plastic ventilated Allentown cages with Beta chip for bedding. Animals were given paper towels and a polyvinylchloride tube for enrichment. The vivarium was temperature and humidity-controlled (21-22°C and 40-60% relative humidity) with standard 12 hr light:12 hr darkness cycles (lights on 08:30-20:30). Sample size was determined based on the pregnancy success and findings of previous work (30,31).

Adult male Long-Evans rats (n=32 total; postnatal day 77-84, 363-435 g; Charles River Laboratories, Kingston, NY, USA) were housed under the same conditions and pair housed with another male. All procedures were approved by the University of British Columbia Animal Care Committee and were in accordance with the Canadian Council on Animal Care. The protocol was not preregistered.

### Diets

After one week of acclimation, all animals were handled daily for 3-4 d. Virgin female rats were randomly assigned to either a CON (1% kcal sucrose; D12450ki; Research Diets Inc., New Brunswick, NJ, USA) or a HSD (26% kcal sucrose; D20061801i; Research Diets, Inc., New Brunswick, NJ, USA) (**Table 1**). The rats were gradually introduced to CON or HSD by adding 6 diet pellets at the bottom of the cage over 3 d. The CON and HSD are isocaloric (3.8 kcal/g) and macro/micro-nutrient matched (10% kcal fat, 20% kcal protein, and 70% kcal carbohydrates). In the HSD, 25% kcal from corn starch is replaced with sucrose. The rats consumed CON or HSD for 19 d of pregnancy.

### Breeding

Females underwent daily vaginal lavage to assess the stage of the estrous cycle. Males were single housed for 2-3 d prior to mating. When a female rat was in proestrus or estrus, the subject was added to a male cage 2-3 hr before lights off. The following morning (within 1 hr of lights on), the female was lavaged to check for sperm. If sperm were present, then the subject was considered pregnant, removed from the male cage, and single housed. The female was monitored for body mass increases to confirm pregnancy. If sperm were absent, then we continued to monitor the female’s body mass and estrous cycle and re-paired the female with the same male when she returned to proestrus or estrus. Each female was mated with a different male.

The day of pregnancy confirmation was defined as E0.5. Upon pregnancy confirmation, females were single housed and fully transitioned onto their respective diets with *ad libitum* access until euthanasia at E19.5. Body mass was measured daily except for the day of euthanasia, and food consumption was measured on E6.5, 12.5, and 19.5. In sum, 30 animals became pregnant and carried fetuses until E19.5 (n=15/diet).

### Tissue collection

Dams were euthanized on E19.5, which is 2-3 d before parturition (36). We selected E19.5 to avoid the increase in glucocorticoid levels close to birth (37), to include the increase in testosterone levels in male fetuses (38), and to match our previous study (30). On E19.5, dams were euthanized between 11:30-13:30 (3-5 hr after lights on). Dams were rapidly and deeply anesthetized using 5% isoflurane and euthanized by rapid decapitation (within 3 min of initial cage disturbance) to minimize the effects of stress on steroid levels (39). For maternal serum, trunk blood was collected and kept on wet ice (30-90 min) and centrifuged for 3-5 min at 5080 g. The serum was immediately frozen on dry ice and stored at -70°C.

Then, the uterine horns were dissected, and the litter size and number of resorptions were recorded. For each fetus, the amniotic fluid was collected using an 18G needle and a syringe. Next, the placenta was removed and rinsed twice in 1× phosphate buffered saline to remove blood. The fetus was decapitated for the brain, and trunk blood was collected using a P20 pipette. The fetal tail was collected to determine sex by genotyping. All samples were immediately frozen on dry ice and stored at -70 °C. One female and one male fetus per dam and their respective placenta and amniotic fluid were used for steroid analysis to control for litter effects (n=12-15/diet/sex).

### Placenta and brain microdissection

Placenta was weighed, coated with Optimal Cutting Temperature compound, and sectioned at 300 µm in a cryostat at -14°C. Each section contained both junctional and labyrinth zones. A 1 mm diameter biopsy punch tool (Miltex, Fisher Scientific) was used to dissect the junctional and labyrinth zones (30,40). Twenty punches were collected per zone (3.66 mg/zone).

The fetal head was also coated with Optimal Cutting Temperature compound and sectioned coronally at 300 µm in a cryostat at -14°C. The NAc, AMY, HYP, vHPC, and VTA were microdissected bilaterally with a 1 mm diameter biopsy punch tool. Depending on the region, a total of 4-8 punches were collected (0.8-1.6 mg/region). The samples were stored in 2 mL polypropylene microcentrifuge tubes containing 5 zirconium ceramic oxide beads (1.4 mm diameter) at -70°C.

### Steroid quantification via LC-MS/MS

Steroids were extracted as previously described (30–32,41). The sample order was randomized and experimenters were blinded. We measured steroids in 5 µL of maternal serum, 10 µL of amniotic fluid, 5 µL of fetal blood, 3.66 mg of each placental zone, and 0.8-1.6 mg of fetal brain. Briefly, for each sample, 1 mL of high-performance liquid chromatography (HPLC)-grade acetonitrile and 20-200 pg of deuterated internal standards (dehydroepiandrosterone (DHEA)-d_6_, progesterone-d_9_, pregnenolone-d_4_, corticosterone-d_8_, testosterone-d_5_, 17β-estradiol-d_4_, aldosterone-d_7_, allopregnanolone-d_4_) were added. Samples were homogenized in a bead homogenizer for 30 s at 4 m/s and then centrifuged for 5 min at 16100 g. Then, 1 mL of supernatant was transferred to pre-cleaned glass culture tubes. Next, 500 µL of HPLC-grade hexane was added, samples were vortexed and centrifuged, and hexane was discarded. Samples were dried in a vacuum centrifuge at 60°C for 45 min. Steroids were resuspended in 55 µL 25% HPLC-grade methanol, centrifuged at 16100 g, and 50 µL of supernatant was transferred to HPLC vials with glass inserts. Samples were stored at -20°C.

A calibration curve from 0.05 to 5000 pg per tube was used for: DHEA, pregnenolone, progesterone, 11-deoxycorticosterone (DOC), corticosterone, DHC, androstenedione, testosterone, estrone, 17β-estradiol, estriol, aldosterone, allopregnanolone, and tetrahydrodeoxycorticosterone. These steroids are important for pregnancy and placental and fetal development (42,43), and their levels are affected by maternal stress and diet (44–46). Duplicates of blanks, double blanks, and quality controls, and triplicates of interassay serum controls (pooled maternal serum) were included.

Steroids were quantified using a SCIEX 6500 QTRAP triple quadrupole tandem mass spectrometer (SCIEX LLC, Framingham, MA). Positive electrospray ionization was used except for estrone, 17β-estradiol, and estriol, which were quantified using negative electrospray ionization. Each analyte was monitored using two multiple reaction monitoring transitions. Internal standards were monitored using one multiple reaction monitoring transition. All blanks, double blanks, and quality controls were acceptable.

### Sucrose quantification *via* LC-MS/MS

Maternal serum (20 µL) was mixed with 160 µL ice-cold methanol and stored at -20°C for 4 hr for protein precipitation and then centrifuged at 14,000 g for 15 min at 4°C. The supernatant was transferred to new tubes and dried at 24°C for 3 hr. Samples were reconstituted in 100 µL acetonitrile/H_2_O (1:1, v/v) on the day of analysis. All solvents were LC-MS grade. A method blank was prepared by subjecting an empty vial to the same extraction procedure. 10 µL aliquots of all maternal serum were combined to create quality control. The sucrose stock solution was prepared at 500 ng/mL in acetonitrile/H_2_O (1:1, v/v) using a pure standard and diluted to 70 ng/mL for retention time confirmation during analysis.

Targeted sucrose quantification was performed on a TSQ Quantis™ Triple Quadrupole mass spectrometer operated in electron spray ionization- and multiple reaction monitoring mode, coupled to a Vanquish™ Horizon ultra-HPLC system (Thermo Fisher Scientific, Mississauga, ON, Canada). The two most intense product fragments were selected for multiple reaction monitoring transitions. Quality control samples were run every six injections, and the standard solution was analyzed at the beginning and end of the run. Further details are in the Supplementary Materials and Methods.

### Data analysis

For steroid analysis, chromatograms were analyzed using MultiQuant Software. If both transitions were absent, then the analyte was considered non-detectable. DHEA, estriol, and tetrahydrodeoxycorticosterone were non-detectable in all samples. Aldosterone and 17β-estradiol were above the lower limit of quantification (LLOQ) in the maternal serum and amniotic fluid only. DOC, corticosterone, DHC, progesterone, and androstenedione were above the LLOQ in all samples. For testosterone, pregnenolone, allopregnanolone, and estrone, at least 50% of samples were above the LLOQ, and for these analytes, values below the LLOQ were imputed using quantile regression imputation of left-censored missing data (47). One dam and the corresponding placenta and fetal samples were excluded from steroid analysis because of complications during euthanasia and excessive handling stress.

For sucrose analysis, raw data were exported as .csv files from chromatograms using Thermo Freestyle. Peak areas and retention times were extracted using a custom Python script. Sucrose identity was confirmed by retention time matching with the standard.

Statistical analysis was conducted on GraphPad Prism 10.4.0, and α was set at ≤ 0.05. If the data did not meet the normality or homogeneity of variance assumptions, then the data were log-transformed before analyses. All data are presented using non-transformed data (mean ± standard mean error). For maternal and pregnancy outcomes and maternal serum analysis (steroids and sucrose), two tailed Student’s t-tests were conducted. For placenta, fetal blood, fetal brain, and amniotic fluid steroid data, two-way analysis of variance was conducted to assess the main effects of Diet and Sex, and their interaction.

## RESULTS

### Maternal and pregnancy outcomes

Maternal HSD during pregnancy alone did not affect total food intake of dams (**Fig. 2A**; t(28) = 0.07, p = 0.95) and body mass gain of dams (**Fig. 2B**; t(28) = 1.08, p = 0.29). The HSD significantly increased sucrose levels in the maternal serum (**Fig. 2C**; t(28) = 10.32, p < 0.0001).

**Figure 2:**
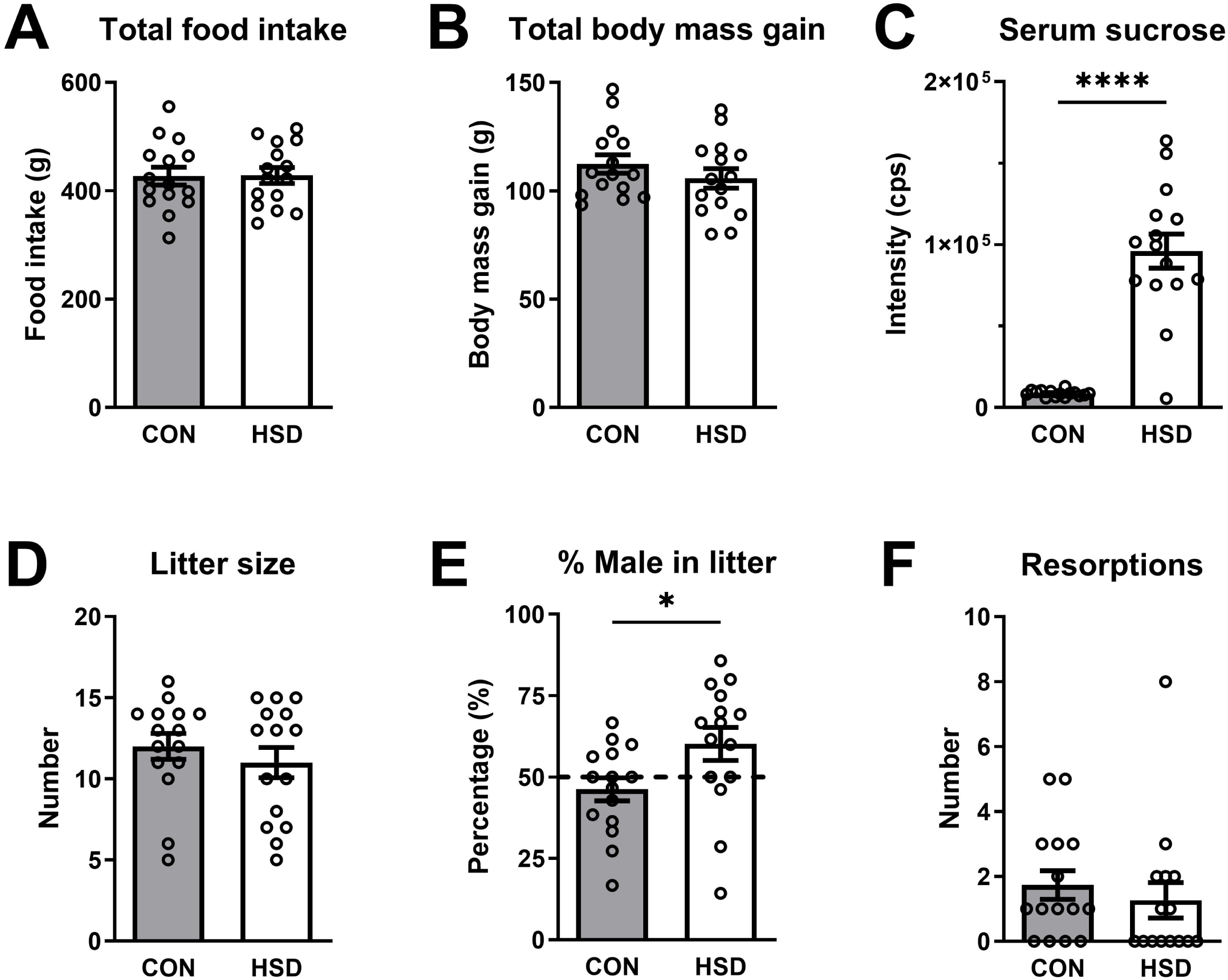
Maternal and pregnancy outcomes. Maternal HSD during pregnancy alone did not alter (**A**) total food intake during pregnancy, (**B**) total body mass gain during pregnancy, but (**C**) significantly increased sucrose levels in the maternal serum. Maternal HSD during pregnancy alone did not change (**D**) litter size and (**F**) number of resorptions, but (**E)** significantly increased the percentage of male offspring per litter. Data are presented as mean ± SEM. n = 15/diet. Data were analyzed using Student’s t-tests. *p ≤ 0.05, ****p < 0.0001. CON: control diet; HSD: high-sucrose diet; cps: counts per second.

Maternal HSD during pregnancy alone did not alter litter size (**Fig. 2D**; t(28) = 0.81, p = 0.42) but significantly increased the percentage of male fetuses in a litter (**Fig. 2E**; t(28) = 2.25, p = 0.03). The number of resorptions did not differ between the diet groups (**Fig. 2F**; t(16) = 0.29, p = 0.78).

### Maternal serum steroids

Maternal HSD during pregnancy alone did not alter levels of DOC, corticosterone, DHC, and aldosterone in the maternal serum (**Fig. 3A-D**; all p values ≥ 0.40). The other steroids measured in the maternal serum were also unaffected by HSD (Supplementary Table 1).

**Figure 3:**
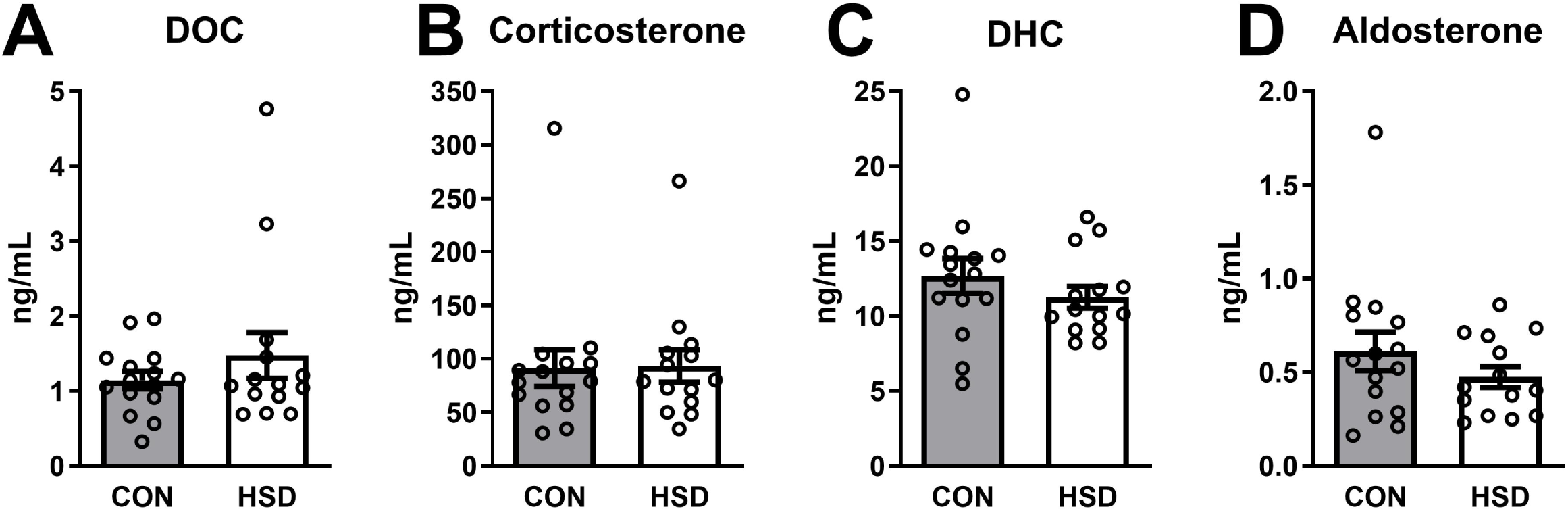
Steroids in the maternal serum. Maternal HSD during pregnancy alone did not alter steroids in maternal serum, including (**A**) DOC, (**B**) corticosterone, (**C**) DHC, and (**D**) aldosterone. Data are presented as mean ± SEM. n = 14-15/diet. Data were analyzed using Student’s t-tests. CON: control diet; HSD: high-sucrose diet; DOC: 11-deoxycorticosterone; DHC: 11-dehydrocorticosterone.

### Placental mass and steroids

Maternal HSD during pregnancy alone did not change placenta mass (**Fig. 4A**; Diet: F(1,47) = 0.34, p = 0.54; Sex: F(1,47) = 2.15, p = 0.15; Diet × Sex: F(1,47) = 0.98, p = 0.33). Maternal HSD did not alter androstenedione and testosterone levels in both zones of the placenta (**Fig. 4C-F**; all p values ≥ 0.50). There was a main effect of Sex for androstenedione and testosterone levels in the junctional zone (males > females, both p values ≤ 0.0002), but not in the labyrinth zone (both p values ≥ 0.13), as before (30). Qualitatively, androstenedione levels were higher than testosterone levels in both zones. There were no Diet × Sex interactions for androstenedione and testosterone levels in both zones (both p values ≥ 0.77). Maternal HSD also did not affect other steroids measured in the placenta (Supplementary Tables 2, 3).

**Figure 4:**
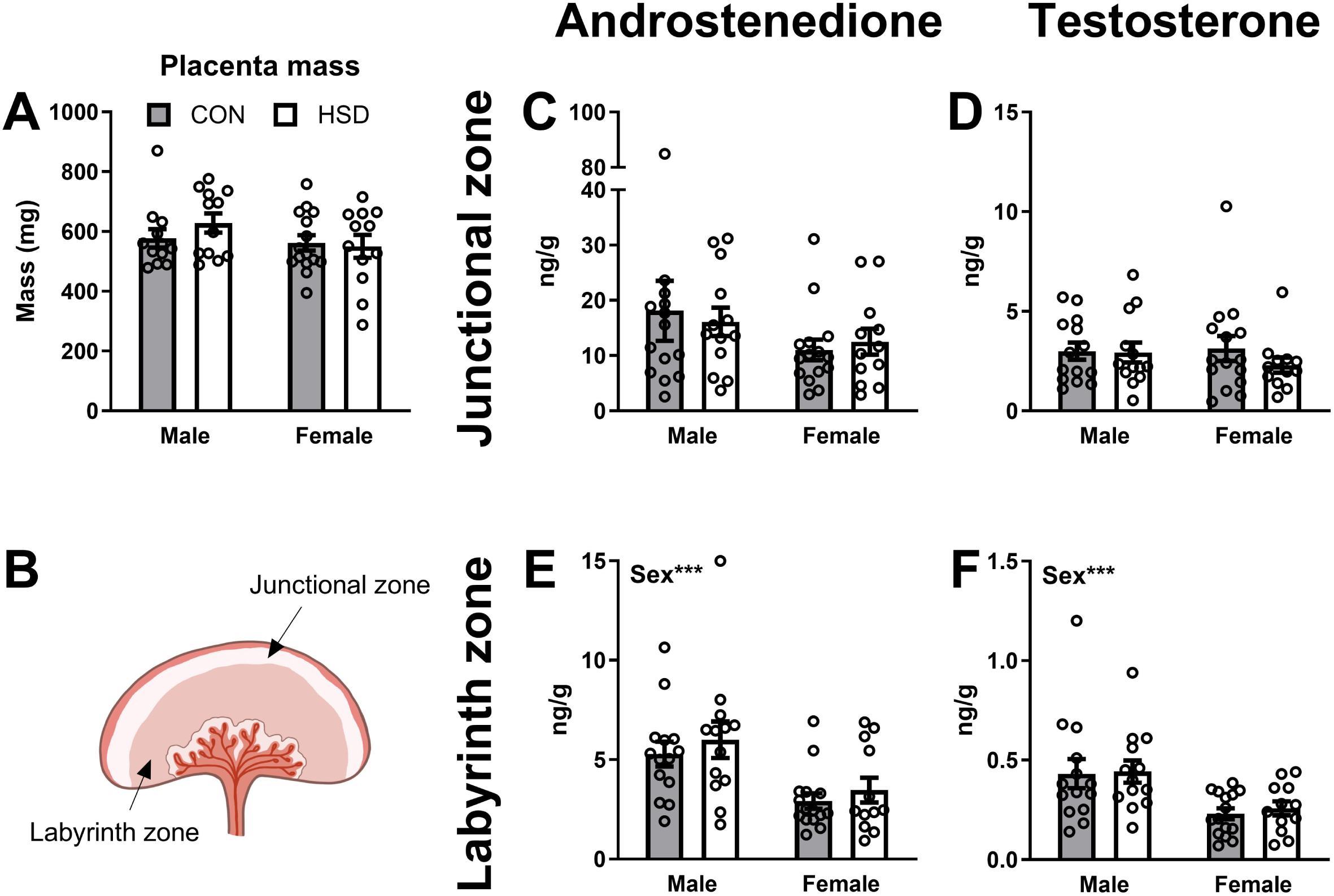
Androgens in the junctional zone and labyrinth zone of the placenta. (**A**) Maternal HSD during pregnancy alone did not alter placenta mass. (**B**) Cross section of the placenta depicting the junctional and labyrinth zones. Maternal HSD during pregnancy alone did not alter (**C,E**) androstenedione and (**D,F**) testosterone levels in the junctional and labyrinth zones of the placenta. Androstenedione and testosterone levels were higher in males than females in the labyrinth zone. Data are presented as mean ± SEM. n = 12-15/diet/sex. Data were analyzed using 2-way ANOVAs. ***p ≤ 0.001. CON: control diet; HSD: high-sucrose diet; ANOVA: analysis of variance.

### Androgens and allopregnanolone in the fetal blood and brain

In the fetal blood, there were no main effects of Diet on androstenedione, testosterone, or allopregnanolone (**Fig. 5A-C**; all p values ≥ 0.41). In the NAc, maternal HSD significantly increased testosterone levels (**Fig. 5E**; Diet: F(1,53) = 4.63, p = 0.04), but did not significantly alter androstenedione and allopregnanolone levels (**Fig. 5D,F**; both p values ≥ 0.12). In the AMY, maternal HSD significantly reduced allopregnanolone (**Fig. 5I**; Diet: F(1,53) = 6.66 p = 0.01) but had no significant effects on androstenedione and testosterone (**Fig. G,H**; both p values p ≥ 0.13). Maternal diet did not affect other steroids measured in the fetal blood and brain (Supplementary Tables 4, 5).

**Figure 5:**
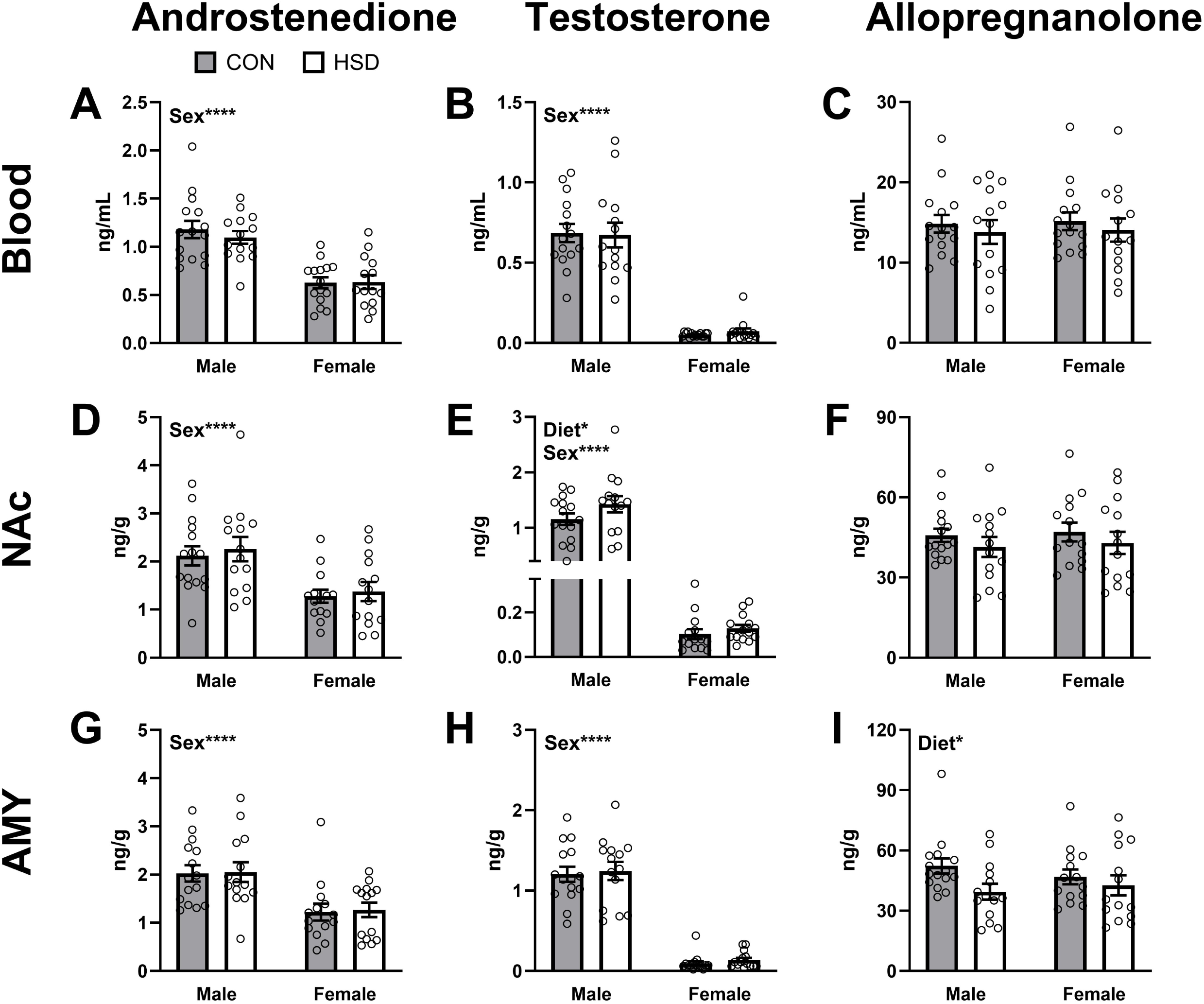
Steroids in the fetal blood and brain. Maternal HSD during pregnancy alone did not affect (**A**) androstenedione, (**B**) testosterone, and (**C**) allopregnanolone in the fetal blood. Maternal HSD during pregnancy alone did not alter (**D**) androstenedione and (**F**) allopregnanolone but significantly increased (**E**) testosterone in the NAc. Maternal HSD during pregnancy alone did not change (**G**) androstenedione and (**H**) testosterone but significantly decreased (**I**) allopregnanolone in the AMY. Males have higher androstenedione and testosterone levels than females in the blood, NAc, and AMY. Data are presented as mean ± SEM. n=14-15/diet/sex. Data were analyzed using 2-way ANOVAs. *p ≤ 0.05; ****p ≤ 0.0001. CON: control diet; HSD: high-sucrose diet; NAc: nucleus accumbens; AMY: amygdala; ANOVA: analysis of variance.

In the fetal blood, NAc, and AMY, there were main effects of Sex for androstenedione and testosterone levels (males > females, all p values < 0.0001). There were no significant Diet × Sex interactions for androstenedione, testosterone, or allopregnanolone (all p values ≥ 0.16). Qualitatively, androstenedione levels were higher than testosterone levels in the fetal blood and brain.

### DHC/corticosterone ratio in the fetal blood and brain

There were no significant main effects of Diet or Diet × Sex interactions on DHC/corticosterone ratio in the fetal blood and fetal brain regions (**Table 2**; all p values ≥ 0.11). There was a significant main effect of Sex in the HYP (**Table 2**; Sex: F(1,53) = 4.40, p = 0.04), but not in other brain regions. In the fetal blood, corticosterone levels were greater than DHC levels (ratio <1). In contrast, in all brain regions, DHC levels were greater than corticosterone levels (ratio >1). Details of the statistical analyses are in Supplementary Table 6.

**Table 2:**
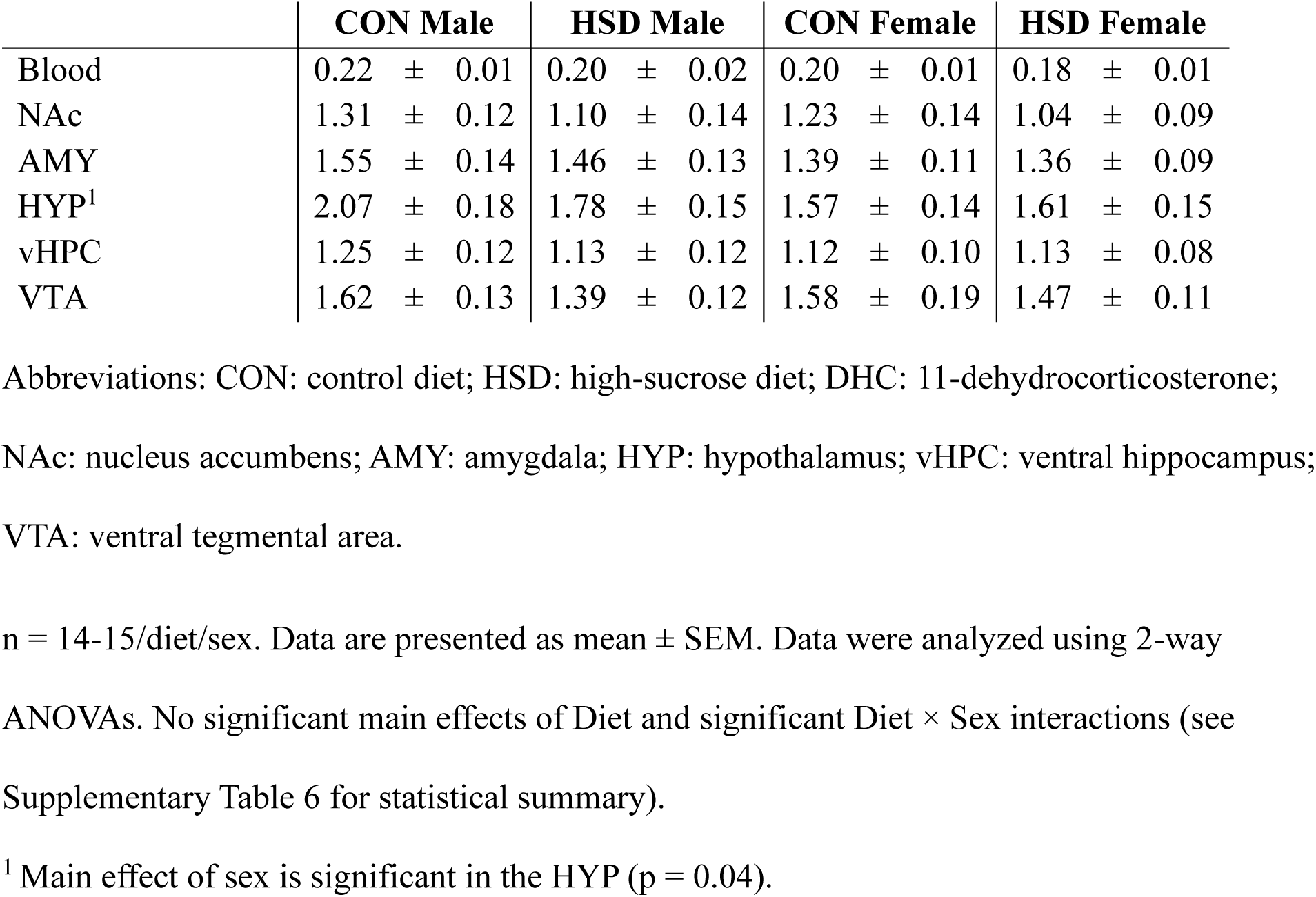
DHC/corticosterone ratio in the blood and brain of fetal offspring exposed to maternal CON or HSD.

### DHC in the amniotic fluid

Maternal HSD significantly reduced DHC levels in the amniotic fluid (**Fig 6C**; Diet: F(1,54) = 4.08, p = 0.05), but did not affect DOC, corticosterone, and aldosterone levels (**Fig 6A,B,D**; all values p ≥ 0.25). There were no significant main effects of Sex or Diet × Sex interactions for DOC, corticosterone, DHC, and aldosterone (all p values ≥ 0.06). Maternal HSD did not affect other steroids measured in the amniotic fluid (Supplementary Tables 7, 8).

**Figure 6:**
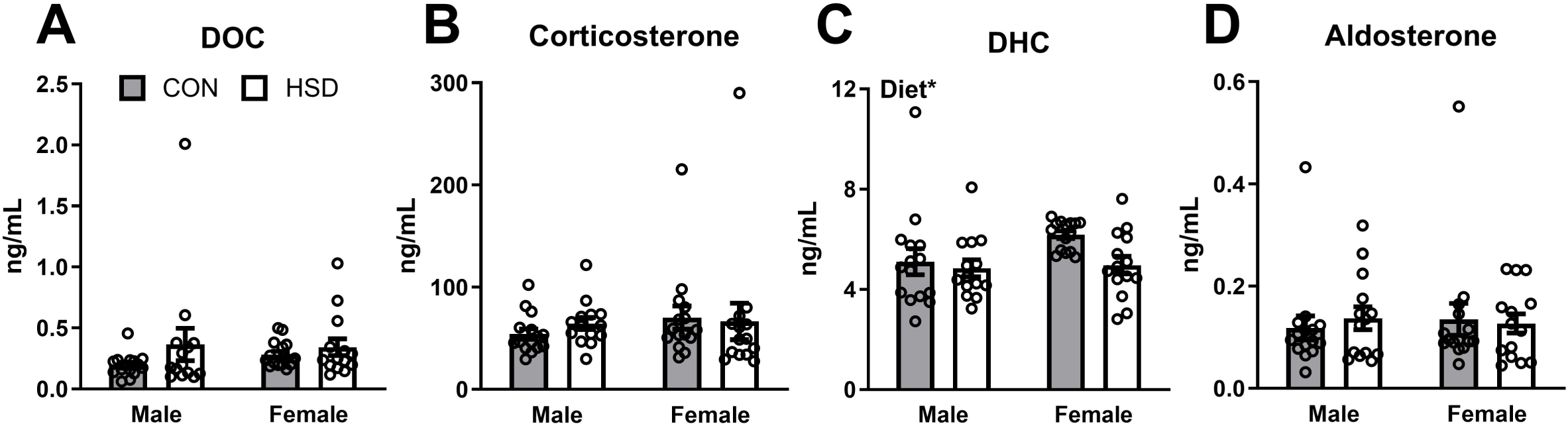
Steroids in the amniotic fluid. Maternal HSD during pregnancy alone did not affect (**A**) DOC, (**B**) corticosterone, and (**D**) aldosterone levels in the amniotic fluid but significantly reduced (**C**) DHC levels in the amniotic fluid. Data are presented as mean ± SEM. n = 14-15/diet/sex. Data were analyzed by 2-way ANOVAs. *p ≤ 0.05. CON: control diet; HSD: high-sucrose diet; DOC: 11-deoxycorticosterone; DHC: 11-dehydrocorticosterone; ANOVA: analysis of variance.

## DISCUSSION

We investigated the effects of a short-term (∼3 wk) maternal HSD during pregnancy alone on maternal, placental, and fetal steroids. Several key results emerge. First, short-term maternal HSD does not alter maternal food intake, maternal body mass, litter size, or resorptions but significantly increases the percentage of males in a litter. Second, maternal, placental, and fetal blood steroid levels, as well as DHC/corticosterone ratio in the fetal blood and brain, are not significantly affected by short-term maternal HSD, in contrast to our previous results with a long-term (∼13 wk) maternal HSD starting 10 wk before pregnancy (30). Third, short-term maternal HSD increases testosterone in the NAc and decreases allopregnanolone in the AMY of the fetal brain. Lastly, short-term maternal HSD decreases DHC in the amniotic fluid. Together, these results suggest that a maternal HSD exclusively during pregnancy is not sufficient to alter maternal, placental, systemic fetal steroids but is sufficient to impact key steroids in critical regions of the fetal brain. In contrast, a long-term maternal HSD has widespread effects on maternal, placental, and fetal steroids (30).

### Maternal HSD during pregnancy alone does not alter maternal food intake and body mass but alters offspring sex ratio

The HSD increases sucrose levels in the maternal serum, which confirms the diet manipulation. Maternal HSD does not affect total food intake and total body mass gain during pregnancy, as before (30,31). Our HSD model is not a maternal obesity model and group differences cannot be attributed to increased maternal caloric intake or adiposity.

Maternal HSD does not change litter size but increases the percentage of males in a litter. The Trivers-Willard hypothesis proposes that better maternal condition leads to a greater proportion of male offspring to maximize maternal reproductive success (48,49). Serum glucose level is a potential indicator of maternal condition and is increased by sucrose intake (50). In *in vitro* studies, higher glucose levels favour the survival of male bovine blastocysts (51). In *in vivo* studies, female voles and mice with higher levels of circulating glucose produce a greater proportion of male offspring in a litter (52,53). Similarly, maternal fructose intake before and during pregnancy increases the proportion of male offspring in rats (54). Thus, maternal sugar intake could alter embryo survival in a sex-specific manner.

### Maternal HSD during pregnancy alone does not impact maternal and placental steroids

Short-term maternal HSD does not alter maternal circulating steroid levels, including glucocorticoids and aldosterone. In contrast, long-term maternal HSD (starting 10 wk before pregnancy) increases DOC and DHC levels, but not corticosterone levels, in maternal serum at E19.5 (30). Other rodent studies have shown decreases in maternal circulating corticosterone levels following 6+ wk of sucrose intake (29,31). Thus, HSD consumption during pregnancy alone is not sufficient to alter maternal serum glucocorticoids.

Maternal HSD during pregnancy alone does not alter placenta mass at E19.5. In contrast, long-term maternal HSD reduces placenta mass at E19.5 (30). Maternal fructose intake during gestation alone decreases female placenta mass at E21.5 in rats (27). Moreover, *in vitro* fertilized embryos that are implanted in mice that consumed 67% fructose for 6 wk have a smaller placenta at E14.5 (55). These findings suggest that a longer duration or higher amount of maternal sugar intake is required to alter placental mass.

Moreover, we do not observe effects of maternal HSD during pregnancy alone on placental steroids. In contrast, our previous results with long-term maternal HSD showed reductions of androstenedione and testosterone in both placenta zones (30). Here, the lack of changes in placental androgens might explain the lack of change in placenta mass, because placental androgens are important for placental development (42).

### Maternal HSD during pregnancy alone increases testosterone in the fetal NAc

The NAc is a key part of the mesocorticolimbic system, an important circuit for executive function and reward processing. The NAc is particularly sensitive to the effects of sugar intake (15,56,57). Moreover, the NAc is modulated by androgens (58), contains androgen receptors (59), and expresses androgenic enzymes (60). Androgen receptor mRNA expression is higher in the NAc than in other regions of the reward circuit, such as the VTA (32). Interestingly, maternal HSD during pregnancy alone increases testosterone levels in the fetal NAc but does not alter testosterone levels in the fetal blood and other regions. In contrast, long-term maternal HSD decreases testosterone levels in the fetal NAc (30) and reduces the expression of *Cyp17a1*, an essential androgenic enzyme, in the NAc of adult male offspring (32). Thus, local testosterone production in the NAc is highly sensitive to maternal HSD in a time-dependent manner. Testosterone levels in the NAc might initially increase following sucrose intake but decrease over time.

Androgens act on the NAc and other nodes of the mesocorticolimbic system to modulate executive function and reward-related behaviours. For example, testosterone administration reduces behavioural flexibility (61), and inhibiting androgen synthesis improves behavioural flexibility (62). Maternal HSD starting 10 wk pre-pregnancy until offspring weaning increases motivation for sugar rewards and increases preference for high-sucrose diet and high-fat diet in adult male offspring (32). These effects of maternal diet on offspring motivation for sugar and on food preference might be mediated by androgen signalling in the NAc.

### Maternal HSD during pregnancy alone decreases allopregnanolone in the fetal AMY

The AMY regulates emotional processing, including fear and anxiety. The short-term maternal HSD reduces allopregnanolone levels in the fetal AMY but does not alter allopregnanolone levels in the fetal blood or other brain regions. The AMY expresses the steroidogenic enzymes to locally synthesize allopregnanolone (63). Allopregnanolone is a positive allosteric modulator of γ-aminobutyric acid (GABA)_A_ receptors, which are present in the AMY and important for inhibitory neurotransmission. Allopregnanolone is also important for the maternal and fetal brain during pregnancy due to its neuroprotective actions (64). In mice, chronic unpredictable stress reduces allopregnanolone levels and reduces mRNA levels of 5α-reductase type I, an enzyme important for allopregnanolone synthesis, in the AMY (65), indicating that allopregnanolone levels in the AMY are sensitive to stress.

Allopregnanolone is implicated in psychiatric disorders, such as anxiety and depression. Low allopregnanolone levels during pregnancy are associated with anxiety and depressive symptoms in pregnant and post-partum women (66,67). Allopregnanolone administration directly into the AMY decreases anxiety-like behaviour (68,69) and decreases depressive-like behaviour in rats (70). Thus, the AMY is a key site of action for the anxiolytic and antidepressant effects of this well-studied neurosteroid.

### Maternal HSD during pregnancy alone does not alter DHC/corticosterone ratio in the fetal blood and brain

Corticosterone can be metabolized into the inactive metabolite DHC via 11β-hydroxysteroid dehydrogenase 2 (11β-HSD2). Here, maternal HSD during pregnancy alone does not significantly affect glucocorticoid levels and the DHC/corticosterone ratio in the fetal blood and brain. In contrast, long-term maternal HSD alters circulating and brain glucocorticoids in the fetal and adult offspring (30,32) and increases the DHC/corticosterone ratio in the fetal brain (30). In mice, maternal sucrose intake starting 4-6 wk before pregnancy and until E18.5 decreases glucocorticoid receptor mRNA in the female fetal brain (29). Maternal fructose or high-fructose corn syrup consumption during pregnancy and lactation increases circulating corticosterone (71,72) and increases 11β-HSD2 mRNA in the hippocampus (24,25) of adult rat male offspring. Overall, long-term maternal sugar intake has prominent effects on glucocorticoid signalling in the offspring.

Females have higher blood corticosterone levels than males at E19.5. A similar sex difference in baseline circulating corticosterone is observed in adult rodents (73), but not in neonatal rodents (74–76). In mice, circulating corticosterone levels are higher in females than males at E18-19 (77,78). This sex difference could be due to greater transport of maternal corticosterone by the female placenta, or differences in glucocorticoid regulation. We do not observe sex differences in placental corticosterone levels, which suggests sex differences in fetal corticosterone production or excretion.

### Maternal HSD during pregnancy alone reduces DHC in the amniotic fluid

Short-term maternal HSD reduces DHC levels in the amniotic fluid. In contrast, long-term maternal HSD increases DHC levels in the amniotic fluid (30), suggesting that the duration of sucrose intake is important. Fetal urine contributes to the amniotic fluid (79) and a reduction in DHC in the amniotic fluid could indicate changes in fetal glucocorticoid excretion. Consistent with the present study, maternal high-fructose corn syrup intake reduces 11β-HSD2 activity in offspring kidneys (71). Moreover, maternal fructose intake reduces 11β-HSD2 expression in the offspring adrenal glands (72). Together, these data suggest that maternal sugar intake alters offspring corticosterone clearance via changes in 11β-HSD2 in the kidneys and adrenal glands.

### Aldosterone levels in the placental and fetal tissues

Aldosterone important for blood pressure regulation, but aldosterone is also involved in the stress response. Circulating aldosterone levels increase following a stressor (35), particularly in the perinatal period (34). High intake of sugars, especially fructose, drives aldosterone production and high blood pressure (80). Long-term maternal HSD increases aldosterone levels in the placenta, fetal blood, fetal brain, and amniotic fluid, suggesting that long-term maternal HSD is a stressor that strongly increases aldosterone levels in the fetus (30).

Short-term maternal HSD does not alter aldosterone levels in maternal serum and amniotic fluid, in contrast to long-term maternal HSD (30). Thus, a longer exposure might be necessary to increase aldosterone levels. Here, aldosterone levels in the placental and fetal tissues were just below the LLOQ. The assay sensitivity in the present study was comparable to that in our previous study (LLOQ = 0.05-0.1 pg) (30), but the background (“noise”) was slightly higher. In our previous study, aldosterone levels were present just above the LLOQ in the CON group and were increased by long-term maternal HSD (30). Here, it is possible that aldosterone levels were unchanged by short-term maternal HSD and just below the LLOQ in both groups.

### Limitations

Rat models allow controlled experiments but have several limitations. First, pregnancy in rats is much shorter (∼21 d) compared to humans, which leads to different neurodevelopmental timeline. The central nervous system of rats is less developed at birth and several neurodevelopmental processes that occur during human pregnancy take place postnatally in rats (81). Second, placental morphology and steroidogenic enzyme expression differ between rats and humans (41,82). For example, CYP17A1 is present in rat, but not human, placenta (82). These differences need to be acknowledged before applying our findings from rat model to human brain health.

## CONCLUSIONS

In conclusion, maternal HSD during pregnancy alone does not alter maternal, placental, and fetal blood steroid levels, but does alter testosterone and allopregnanolone levels in key regions of the fetal brain and does decrease DHC levels in the amniotic fluid. Thus, specific regions of the fetal brain are sensitive to even short-term maternal sucrose intake. In contrast, long-term maternal HSD has widespread effects on maternal, placental, and fetal steroids. Our findings, together with previous animal studies, demonstrate that pregnancy sugar consumption impacts the developing offspring brain. Future studies will examine brain steroidogenic enzymes that produce testosterone and allopregnanolone and examine relevant behaviours, such as reward seeking and anxiety-like behaviours, in offspring.

## ACKNOWLEDGEMENTS

We thank Hui W. Chen for assistance with data collection; Dr. Asmita Poudel for assistance with mass spectrometry; Dr. Angela Devlin for comments on the manuscript; Dr. Melody Salehzadeh for help with Fig. 1; Zeyu Yang for help with Fig 4B; UBC Centre for Disease Modelling staff for animal husbandry; and UBC Genotyping facility for data collection.

## AUTHOR CONTRIBUTIONS

MMJ, DRS, and KKS designed research; MMJ, MI, and YNH conducted research; MMJ and YNH analyzed data; MMJ and KKS wrote the paper; MMJ, TH, and KKS had primary responsibility for final content. All authors read and approved the final manuscript.

## DATA AVAILABILITY

Data described in the manuscript will be made publicly and freely available at: DOI 10.17605/OSF.IO/3PMN4

## LIST OF ABBREVIATIONS

11β-HSD2: 11β-hydroxysteroid dehydrogenase 2
AMY: amygdala CON: control diet
DHC: 11-dehydrocorticosterone
DHEA: dehydroepiandrosterone
DOC: 11-deoxycorticosterone
E: embryonic day
HPLC: high performance liquid chromatography
HSD: high-sucrose diet
HYP: hypothalamus
LLOQ: lower limit of quantification
LC-MS/MS: liquid chromatography-tandem mass spectrometry
NAc: nucleus accumbens
vHPC: ventral hippocampus
VTA: ventral tegmental area

## REFERENCES

1. Walton J, Bell H, Re R, Nugent AP. Current perspectives on global sugar consumption: definitions, recommendations, population intakes, challenges and future direction. Nutr Res Rev 2023;36:1–22.

2. Cioffi CE, Figueroa J, Welsh JA. Added Sugar Intake among Pregnant Women in the United States: National Health and Nutrition Examination Survey 2003-2012. J Acad Nutr Diet 2018;118:886–895.e1.

3. Crozier SR, Robinson SM, Godfrey KM, Cooper C, Inskip HM. Women’s Dietary Patterns Change Little from Before to During Pregnancy. J Nutr 2009;139:1956–63.

4. Goran MI, Plows JF, Ventura EE. Effects of consuming sugars and alternative sweeteners during pregnancy on maternal and child health: evidence for a secondhand sugar effect. Proc Nutr Soc Cambridge University Press; 2019;78:262–71.

5. Jen V, Erler NS, Tielemans MJ, Braun KV, Jaddoe VW, Franco OH, Voortman T. Mothers’ intake of sugar-containing beverages during pregnancy and body composition of their children during childhood: the Generation R Study1, 2, 3. Am J Clin Nutr 2017;105:834–41.

6. Murrin C, Shrivastava A, Kelleher CC. Maternal macronutrient intake during pregnancy and 5 years postpartum and associations with child weight status aged five. Eur J Clin Nutr 2013;67:670–9.

7. Gracner T, Boone C, Gertler PJ. Exposure to sugar rationing in the first 1000 days of life protected against chronic disease. Science 2024;0:eadn5421.

8. Johnson RK, Appel LJ, Brands M, Howard BV, Lefevre M, Lustig RH, Sacks F, Steffen LM, Wylie-Rosett J. Dietary Sugars Intake and Cardiovascular Health. Circulation 2009;120:1011–20.

9. Stanhope KL. Sugar consumption, metabolic disease and obesity: The state of the controversy. Crit Rev Clin Lab Sci 2016;53:52–67.

10. Jacques A, Chaaya N, Beecher K, Ali SA, Belmer A, Bartlett S. The impact of sugar consumption on stress driven, emotional and addictive behaviors. Neurosci Biobehav Rev 2019;103:178–99.

11. Witek K, Wydra K, Filip M. A High-Sugar Diet Consumption, Metabolism and Health Impacts with a Focus on the Development of Substance Use Disorder: A Narrative Review. Nutrients 2022;14:2940.

12. Beecher K, Alvarez Cooper I, Wang J, Walters SB, Chehrehasa F, Bartlett SE, Belmer A. Long-Term Overconsumption of Sugar Starting at Adolescence Produces Persistent Hyperactivity and Neurocognitive Deficits in Adulthood. Front Neurosci Frontiers; 2021;15.

13. Beecher K, Wang J, Jacques A, Chaaya N, Chehrehasa F, Belmer A, Bartlett SE. Sucrose Consumption Alters Serotonin/Glutamate Co-localisation Within the Prefrontal Cortex and Hippocampus of Mice. Front Mol Neurosci Frontiers; 2021;14.

14. Hamelin H, Poizat G, Florian C, Kursa MB, Pittaras E, Callebert J, Rampon C, Taouis M, Hamed A, Granon S. Prolonged Consumption of Sweetened Beverages Lastingly Deteriorates Cognitive Functions and Reward Processing in Mice. Cereb Cortex 2022;32:1365–78.

15. Tukey DS, Ferreira JM, Antoine SO, D’amour JA, Ninan I, Cabeza De Vaca S, Incontro S, Wincott C, Horwitz JK, Hartner DT, et al. Sucrose Ingestion Induces Rapid AMPA Receptor Trafficking. J Neurosci 2013;33:6123–32.

16. Choi CS, Kim P, Park JH, Gonzales ELT, Kim KC, Cho KS, Ko MJ, Yang SM, Seung H, Han S-H, et al. High sucrose consumption during pregnancy induced ADHD-like behavioral phenotypes in mice offspring. J Nutr Biochem 2015;26:1520–6.

17. Mizera J, Kazek G, Niedzielska-Andres E, Pomierny-Chamiolo L. Maternal high-sugar diet results in NMDA receptors abnormalities and cognitive impairment in rat offspring. FASEB J 2021;35:e21547.

18. Kuang H, Sun M, Lv J, Li J, Wu C, Chen N, Bo L, Wei X, Gu X, Liu Z, et al. Hippocampal apoptosis involved in learning deficits in the offspring exposed to maternal high sucrose diets. J Nutr Biochem 2014;25:985–90.

19. He A, Zhang Y, Yang Y, Li L, Feng X, Wei B, Zhu D, Liu Y, Wu L, Zhang L, et al. Prenatal high sucrose intake affected learning and memory of aged rat offspring with abnormal oxidative stress and NMDARs/Wnt signaling in the hippocampus. Brain Res 2017;1669:114–21.

20. Bukhari SHF, Clark OE, Williamson LL. Maternal high fructose diet and neonatal immune challenge alter offspring anxiety-like behavior and inflammation across the lifespan. Life Sci 2018;197:114–21.

21. Chakraborti A, Graham C, Chehade S, Vashi B, Umfress A, Kurup P, Vickers B, Chen HA, Telange R, Berryhill T, et al. High Fructose Corn Syrup-Moderate Fat Diet Potentiates Anxio-Depressive Behavior and Alters Ventral Striatal Neuronal Signaling. Front Neurosci 2021;15:669410.

22. Liu W-C, Wu C-W, Fu M-H, Tain Y-L, Liang C-K, Hung C-Y, Chen I-C, Lee Y-C, Wu C-Y, Wu KLH. Maternal high fructose-induced hippocampal neuroinflammation in the adult female offspring via PPARγ-NF-κB signaling. J Nutr Biochem 2020;81:108378.

23. Davis MR, Hester KK, Shawron KM, Lucas EA, Smith BJ, Clarke SL. Comparisons of the iron deficient metabolic response in rats fed either an AIN-76 or AIN-93 based diet. Nutr Metab 2012;9:95.

24. Mizuno G, Munetsuna E, Yamada H, Ando Y, Yamazaki M, Murase Y, Kondo K, Ishikawa H, Teradaira R, Suzuki K, et al. Fructose intake during gestation and lactation differentially affects the expression of hippocampal neurosteroidogenic enzymes in rat offspring. Endocr Res 2017;42:71–7.

25. Ohashi K, Ando Y, Munetsuna E, Yamada H, Yamazaki M, Nagura A, Taromaru N, Ishikawa H, Suzuki K, Teradaira R. Maternal fructose consumption alters messenger RNA expression of hippocampal StAR, PBR, P450(11β), 11β-HSD, and 17β-HSD in rat offspring. Nutr Res 2015;35:259–64.

26. Rey M, Coirini H, Marchena A, González Deniselle MC, Kruse MS. Effects of metformin on behavioral alterations produced by chronic sucrose consumption in male rats. J Neuroendocrinol 2024;36:e13362.

27. Vickers MH, Clayton ZE, Yap C, Sloboda DM. Maternal Fructose Intake during Pregnancy and Lactation Alters Placental Growth and Leads to Sex-Specific Changes in Fetal and Neonatal Endocrine Function. Endocrinology 2011;152:1378–87.

28. Hirai S, Miwa H, Tanaka T, Toriumi K, Kunii Y, Shimbo H, Sakamoto T, Hino M, Izumi R, Nagaoka A, et al. High-sucrose diets contribute to brain angiopathy with impaired glucose uptake and psychosis-related higher brain dysfunctions in mice. Sci Adv 2021;7:eabl6077.

29. Kulhanek D, Rao RB, Paulsen ME. Excess sucrose intake during pregnancy programs fetal brain glucocorticoid receptor expression in female but not male C57Bl/6J mice. Obes Sci Pract 2021;7:462–72.

30. Seib DR, Jung MM, Soma KK. Maternal sucrose consumption alters steroid levels in the mother, placenta and fetus. J Endocrinol 2025;265:e240238.

31. Tobiansky DJ, Kachkovski GV, Enos RT, Schmidt KL, Murphy EA, Soma KK. Sucrose consumption alters steroid and dopamine signalling in the female rat brain. J Endocrinol 2020;245:231–46.

32. Tobiansky DJ, Kachkovski GV, Enos RT, Schmidt KL, Murphy EA, Floresco SB, Soma KK. Maternal sucrose consumption alters behaviour and steroids in adult rat offspring. J Endocrinol 2021;251:161–80.

33. Yang Q, Zhang Z, Gregg EW, Flanders WD, Merritt R, Hu FB. Added Sugar Intake and Cardiovascular Diseases Mortality Among US Adults. JAMA Intern Med 2014;174:516.

34. Varga J, Ferenczi S, Kovács KJ, Garafova A, Jezova D, Zelena D. Comparison of Stress-Induced Changes in Adults and Pups: Is Aldosterone the Main Adrenocortical Stress Hormone during the Perinatal Period in Rats? PLOS ONE 2013;8:e72313.

35. Sánchez-Solís CN, Cuevas Romero E, Soto-Rodríguez I, De Lourdes Arteaga-Castañeda M, De León-Ramírez YM, Rodríguez-Antolín J, Nicolás-Toledo L. High-sucrose diet potentiates hyperaldosteronism and renal injury induced by stress in young adult rats. Clin Exp Pharmacol Physiol 2020;47:1985–94.

36. Modlinska K, Pisula W. The Norway rat, from an obnoxious pest to a laboratory pet. eLife 2020;9:e50651.

37. Voogt J, Sar M, Meites J. Influence of cycling, pregnancy, labor, and suckling on corticosterone-ACTH levels. Am J Physiol-Leg Content 1969;216:655–8.

38. Ward IL, Ward OB, Affuso JD, Long WD, French JA, Hendricks SE. Fetal testosterone surge: specific modulations induced in male rats by maternal stress and/or alcohol consumption. Horm Behav 2003;43:531–9.

39. Taves MD, Ma C, Heimovics SA, Saldanha CJ, Soma KK. Measurement of Steroid Concentrations in Brain Tissue: Methodological Considerations. Front Endocrinol 2011;2:39.

40. Charest PL, Vrolyk V, Herst P, Lessard M, Sloboda DM, Dalvai M, Haruna J, Bailey JL, Benoit-Biancamano M-O. Histomorphologic Analysis of the Late-term Rat Fetus and Placenta. Toxicol Pathol 2018;46:158–68.

41. Price HR, Jalabert C, Seib DR, Ma C, Lai D, Soma KK, Collier AC. Measurement of Steroids in the Placenta, Maternal Serum, and Fetal Serum in Humans, Rats, and Mice: A Technical Note. Separations 2023;10:221.

42. Parsons AM, Bouma GJ. A Potential Role and Contribution of Androgens in Placental Development and Pregnancy. Life 2021;11:644.

43. Noyola-Martínez N, Halhali A, Barrera D. Steroid hormones and pregnancy. Gynecol Endocrinol 2019;35:376–84.

44. Brunton PJ. Programming the Brain and Behaviour by Early-Life Stress: A Focus on Neuroactive Steroids. J Neuroendocrinol 2015;27:468–80.

45. Sze Y, Brunton PJ. How is prenatal stress transmitted from the mother to the fetus? J Exp Biol 2024;227:jeb246073.

46. Bolton JL, and Bilbo SD. Developmental programming of brain and behavior by perinatal diet: focus on inflammatory mechanisms. Dialogues Clin Neurosci Taylor & Francis; 2014;16:307–20.

47. Wei R, Wang J, Jia E, Chen T, Ni Y, Jia W. GSimp: A Gibbs sampler based left-censored missing value imputation approach for metabolomics studies. PLOS Comput Biol 2018;14:e1005973.

48. Douhard M. Offspring sex ratio in mammals and the Trivers-Willard hypothesis: In pursuit of unambiguous evidence. BioEssays 2017;39:1700043.

49. Trivers RL, Willard DE. Natural Selection of Parental Ability to Vary the Sex Ratio of Offspring. Science 1973;179:90–2.

50. Ozkan H, Topsakal S, Ozmen O. Investigation of the diabetic effects of maternal high-glucose diet on rats. Biomed Pharmacother 2019;110:609–17.

51. Kimura K, Spate LD, Green MP, Roberts RM. Effects of D-glucose concentration, D-fructose, and inhibitors of enzymes of the pentose phosphate pathway on the development and sex ratio of bovine blastocysts. Mol Reprod Dev 2005;72:201–7.

52. Helle S, Laaksonen T, Adamsson A, Paranko J, Huitu O. Female field voles with high testosterone and glucose levels produce male-biased litters. Anim Behav 2008;75:1031–9.

53. Machado AF, Zimmerman EF, Hovland Jr. DN, Weiss R, Collins MD. Diabetic Embryopathy in C57BL/6J Mice: Altered Fetal Sex Ratio and Impact of the Splotch Allele. Diabetes 2001;50:1193–9.

54. Gray C, Long S, Green C, Gardiner SM, Craigon J, Gardner DS. Maternal Fructose and/or Salt Intake and Reproductive Outcome in the Rat: Effects on Growth, Fertility, Sex Ratio, and Birth Order. Biol Reprod 2013;89:51, 1–8.

55. Saben JL, Asghar Z, Rhee JS, Drury A, Scheaffer S, Moley KH. Excess Maternal Fructose Consumption Increases Fetal Loss and Impairs Endometrial Decidualization in Mice. Endocrinology 2016;157:956–68.

56. Hajnal A, Smith GP, Norgren R. Oral sucrose stimulation increases accumbens dopamine in the rat. Am J Physiol-Regul Integr Comp Physiol 2004;286:R31–7.

57. Rada P, Avena NM, Hoebel BG. Daily bingeing on sugar repeatedly releases dopamine in the accumbens shell. Neuroscience 2005;134:737–44.

58. Tobiansky DJ, Wallin-Miller KG, Floresco SB, Wood RI, Soma KK. Androgen Regulation of the Mesocorticolimbic System and Executive Function. Front Endocrinol 2018;9:279.

59. Low KL, Tomm RJ, Ma C, Tobiansky DJ, Floresco SB, Soma KK. Effects of aging on testosterone and androgen receptors in the mesocorticolimbic system of male rats. Horm Behav 2020;120:104689.

60. Seib DR, Tobiansky DJ, Meitzen J, Floresco SB, Soma KK. Neurosteroids and the mesocorticolimbic system. Neurosci Biobehav Rev 2023;153:105356.

61. Wallin KG, Wood RI. Anabolic–androgenic steroids impair set-shifting and reversal learning in male rats. Eur Neuropsychopharmacol 2015;25:583–90.

62. Tomm RJ, Seib DR, Kachkovski GV, Schweitzer HR, Tobiansky DJ, Floresco SB, Soma KK. Androgen synthesis inhibition increases behavioural flexibility and mPFC tyrosine hydroxylase in gonadectomized male rats. J Neuroendocrinol 2022;34:e13128.

63. Agís-Balboa RC, Pinna G, Zhubi A, Maloku E, Veldic M, Costa E, Guidotti A. Characterization of brain neurons that express enzymes mediating neurosteroid biosynthesis. Proc Natl Acad Sci 2006;103:14602–7.

64. Brunton PJ, Russell JA, Hirst JJ. Allopregnanolone in the brain: Protecting pregnancy and birth outcomes. Prog Neurobiol 2014;113:106–36.

65. Walton NL, Antonoudiou P, Barros L, Dargan T, DiLeo A, Evans-Strong A, Gabby J, Howard S, Paracha R, Sánchez EJ, et al. Impaired Endogenous Neurosteroid Signaling Contributes to Behavioral Deficits Associated With Chronic Stress. Biol Psychiatry 2023;94:249–61.

66. Hellgren C, Åkerud H, Skalkidou A, Bäckström T, Sundström-Poromaa I. Low Serum Allopregnanolone Is Associated with Symptoms of Depression in Late Pregnancy. Neuropsychobiology 2014;69:147–53.

67. Osborne LM, Betz JF, Yenokyan G, Standeven LR, Payne JL. The Role of Allopregnanolone in Pregnancy in Predicting Postpartum Anxiety Symptoms. Front Psychol 2019;10.

68. Akwa Y, Purdy RH, Koob GF, Britton. The amygdala mediates the anxiolytic-like effect of the neurosteroid allopregnanolone in rat. Behav Brain Res 1999;106:119–25.

69. Engin E, Treit D. The anxiolytic-like effects of allopregnanolone vary as a function of intracerebral microinfusion site: the amygdala, medial prefrontal cortex, or hippocampus. Behav Pharmacol 2007;18:461–70.

70. Shirayama Y, Muneoka K, Fukumoto M, Tadokoro S, Fukami G, Hashimoto K, Iyo M. Infusions of allopregnanolone into the hippocampus and amygdala, but not into the nucleus accumbens and medial prefrontal cortex, produce antidepressant effects on the learned helplessness rats. Hippocampus 2011;21:1105–13.

71. Nouchi Y, Munetsuna E, Yamada H, Yamazaki M, Ando Y, Mizuno G, Ikeya M, Kageyama I, Wakasugi T, Teshigawara A, et al. Maternal High-Fructose Corn Syrup Intake Impairs Corticosterone Clearance by Reducing Renal 11β-Hsd2 Activity via miR-27a-Mediated Mechanism in Rat Offspring. Nutrients 2023;15:2122.

72. Munetsuna E, Yamada H, Yamazaki M, Ando Y, Mizuno G, Hattori Y, Sadamoto N, Ishikawa H, Ohta Y, Fujii R, et al. Maternal high-fructose intake increases circulating corticosterone levels via decreased adrenal corticosterone clearance in adult offspring. J Nutr Biochem 2019;67:44–50.

73. Belz EE, Kennell JS, Czambel RK, Rubin RT, Rhodes ME. Environmental enrichment lowers stress-responsive hormones in singly housed male and female rats. Pharmacol Biochem Behav 2003;76:481–6.

74. Hamden JE, Gray KM, Salehzadeh M, Soma KK. Isoflurane stress induces region-specific glucocorticoid levels in neonatal mouse brain. J Endocrinol 2022;255:61–74.

75. Hamden JE, Salehzadeh M, Bajaj H, Li MX, Soma KK. Lipopolysaccharide differentially alters systemic and brain glucocorticoid levels in neonatal and adult mice. J Neuroendocrinol 2024;e13481.

76. Salehzadeh M, Hamden JE, Li MX, Bajaj H, Wu RS, Soma KK. Glucocorticoid Production in Lymphoid Organs: Acute Effects of Lipopolysaccharide in Neonatal and Adult Mice. Endocrinology 2022;163:bqab244.

77. Montano MM, Wang M-H, Even MD, Vom Saal FS. Serum corticosterone in fetal mice: Sex differences, circadian changes, and effect of maternal stress. Physiol Behav 1991;50:323–9.

78. Montano MM, Wang M-H, Vom Saal FS. Sex differences in plasma corticosterone in mouse fetuses are mediated by differential placental transport from the mother and eliminated by maternal adrenalectomy or stress. J Reprod Fertil 1993;99:283–90.

79. Underwood MA, Gilbert WM, Sherman MP. Amniotic Fluid: Not Just Fetal Urine Anymore. J Perinatol 2005;25:341–8.

80. Hahn K, Rodriguez-Iturbe B, Winterberg B, Sanchez-Lozada LG, Kanbay M, Lanaspa MA, Johnson RJ. Primary aldosteronism: A consequence of sugar and western Diet? Med Hypotheses 2022;160:110796.

81. Zeiss CJ. Comparative Milestones in Rodent and Human Postnatal Central Nervous System Development. Toxicol Pathol 2021;49:1368–73.

82. Furukawa S, Tsuji N, Sugiyama A. Morphology and physiology of rat placenta for toxicological evaluation. J Toxicol Pathol 2019;32:1–17.

