## Supplementary Materials for "Maternal sucrose intake during pregnancy alone disrupts testosterone and allopregnanolone levels in the fetal brain of rats"

**Supplementary Materials and Methods: LC-MS/MS parameters for sucrose quantification**

Hydrophilic interaction chromatography (HILIC) was carried out using a Waters Atlantis™ Premier BEH Z-HILIC VanGuard™ Fit column (2.5 µm, 2.1 mm × 150 mm, 95 Å) (Milford, MA) maintained at 35°C. Mobile phase A was acetonitrile/$H_2O$ (5:95, v/v) with 10 mM $NH_4Ac$, adjusted to pH 9.8 using ammonium hydroxide. Mobile phase B was acetonitrile/$H_2O$ (95:5, v/v). The liquid chromatography gradient was as follows: 0 min, 95% B; 20 min, 5% B; 21 min, 95% B; 35 min, 95% B. The flow rate was 0.3 mL/min, and injection volume was 5 µL. The electrospray ionization source was operated with a vaporizer temperature of 350°C and capillary voltage of 2500 V. Collision energy and RF lens voltages were optimized for the multiple reaction monitoring transitions. The two monitored transitions were as follows (precursor m/z → product m/z): 340.992 → 88.867 and 340.992 → 187.967. The pure standard retention time was 7.01 min and the sample retention time was 7.04 min.

**Supplementary Table 1: All steroids measured in the maternal serum (ng/mL).**

| | CON | | | HSD | | | t | p |
|---|---|---|---|---|---|---|---|---|
| Pregnenolone | 52.60 | ± | 11.65 | 46.38 | ± | 7.33 | 0.17 | 0.86 |
| Progesterone | 124.40 | ± | 6.35 | 116.40 | ± | 3.35 | 1.10 | 0.28 |
| DOC | 1.14 | ± | 0.12 | 1.48 | ± | 0.31 | 0.85 | 0.40 |
| Corticosterone | 91.33 | ± | 17.14 | 93.38 | ± | 15.14 | 0.26 | 0.80 |
| DHC | 12.68 | ± | 1.15 | 11.25 | ± | 0.73 | 0.76 | 0.46 |
| Androstenedione | 0.57 | ± | 0.04 | 0.57 | ± | 0.04 | 0.13 | 0.90 |
| Testosterone | 0.25 | ± | 0.05 | 0.18 | ± | 0.02 | 1.28 | 0.21 |
| Allopregnanolone | 15.65 | ± | 1.09 | 14.27 | ± | 1.28 | 0.83 | 0.41 |
| Aldosterone | 0.61 | ± | 0.10 | 0.48 | ± | 0.06 | 0.81 | 0.43 |
| Estrone | 0.05 | ± | 0.00 | 0.05 | ± | 0.01 | 0.22 | 0.83 |
| 17β-estradiol | 0.09 | ± | 0.01 | 0.08 | ± | 0.01 | 0.76 | 0.46 |

Abbreviations: CON: control diet; HSD: high-sucrose diet; DOC: 11-deoxycorticosterone; DHC: 11-dehydrocorticosterone.

n = 14-15/diet. Data are presented as mean ± SEM. Data were analyzed using t-tests.

**Supplementary Table 2: Additional steroids measured in the placenta (ng/g).**

| | CON Male | HSD Male | CON Female | HSD Female |
|---|---|---|---|---|
| **Pregnenolone** | | | | |
| Junctional zone | 55.07 ± 7.01 | 71.01 ± 7.47 | 64.54 ± 10.52 | 67.86 ± 10.90 |
| Labyrinth zone | 38.57 ± 6.07 | 50.02 ± 6.44 | 42.29 ± 4.71 | 47.33 ± 4.48 |
| **Progesterone** | | | | |
| Junctional zone | 45.99 ± 5.70 | 37.14 ± 2.85 | 42.28 ± 3.01 | 44.00 ± 5.39 |
| Labyrinth zone | 30.86 ± 4.13 | 33.99 ± 4.60 | 38.56 ± 4.40 | 36.03 ± 4.37 |
| **DOC** | | | | |
| Junctional zone | 5.33 ± 0.54 | 7.29 ± 2.15 | 5.53 ± 0.49 | 5.12 ± 0.44 |
| Labyrinth zone | 6.65 ± 0.92 | 9.80 ± 2.79 | 7.29 ± 0.74 | 6.70 ± 0.43 |
| **Corticosterone** | | | | |
| Junctional zone | 46.80 ± 4.41 | 46.15 ± 8.20 | 38.91 ± 3.45 | 45.68 ± 4.78 |
| Labyrinth zone | 63.15 ± 4.96 | 73.72 ± 9.25 | 64.60 ± 4.55 | 72.46 ± 5.75 |
| **DHC** | | | | |
| Junctional zone | 127.56 ± 8.22 | 110.45 ± 8.52 | 125.12 ± 9.45 | 127.64 ± 10.28 |
| Labyrinth zone | 148.91 ± 8.94 | 143.52 ± 9.72 | 159.04 ± 10.36 | 147.06 ± 9.20 |
| **Allopregnanolone** | | | | |
| Junctional zone | 100.15 ± 8.91 | 84.79 ± 9.38 | 86.38 ± 7.44 | 83.30 ± 9.15 |
| Labyrinth zone | 88.80 ± 6.81 | 81.99 ± 6.50 | 79.88 ± 6.66 | 78.07 ± 7.14 |
| **Estrone** | | | | |
| Junctional zone | 0.06 ± 0.01 | 0.06 ± 0.01 | 0.06 ± 0.01 | 0.07 ± 0.01 |
| Labyrinth zone | 0.35 ± 0.03 | 0.35 ± 0.03 | 0.34 ± 0.02 | 0.38 ± 0.03 |

Abbreviations: CON: control diet; HSD: high-sucrose diet; DOC: 11-deoxycorticosterone; DHC: 11-dehydrocorticosterone.

n = 12-15/diet/sex. Data are presented as mean ± SEM. Data were analyzed using 2-way ANOVAs. No significant main effects of Diet or Sex or interactions; see Supplementary Table 3 for statistical summary. Aldosterone and 17β-estradiol were below the lower limit of quantification.

**Supplementary Table 3: Statistical summary for all steroids measured in the placenta.**

| | Diet | | Sex | | Diet × Sex | |
|---|---|---|---|---|---|---|
| | **F** | **p** | **F** | **p** | **F** | **p** |
| **Pregnenolone** | | | | | | |
| Junctional zone | 1.92 | 0.17 | 0.01 | 0.93 | 0.47 | 0.50 |
| Labyrinth zone | 3.33 | 0.07 | 0.32 | 0.57 | 0.39 | 0.54 |
| **Progesterone** | | | | | | |
| Junctional zone | 0.81 | 0.37 | 0.27 | 0.60 | 0.72 | 0.40 |
| Labyrinth zone | <0.0001 | 0.99 | 1.61 | 0.21 | 0.38 | 0.54 |
| **DOC** | | | | | | |
| Junctional zone | 0.02 | 0.89 | 0.04 | 0.83 | 0.41 | 0.52 |
| Labyrinth zone | 0.14 | 0.71 | 0.08 | 0.77 | 0.41 | 0.52 |
| **Corticosterone** | | | | | | |
| Junctional zone | 0.02 | 0.90 | 0.17 | 0.69 | 1.42 | 0.24 |
| Labyrinth zone | 2.15 | 0.15 | 0.00 | 0.99 | 0.05 | 0.83 |
| **DHC** | | | | | | |
| Junctional zone | 0.63 | 0.43 | 0.65 | 0.42 | 1.15 | 0.29 |
| Labyrinth zone | 0.80 | 0.37 | 0.50 | 0.48 | 0.12 | 0.74 |
| **Androstenedione** | | | | | | |
| Junctional zone | 0.13 | 0.72 | 2.39 | 0.13 | 0.01 | 0.93 |
| Labyrinth zone | 0.33 | 0.57 | 17.43 | **0.0001** | <0.0001 | 0.99 |
| **Testosterone** | | | | | | |
| Junctional zone | 0.46 | 0.50 | 0.60 | 0.44 | 0.09 | 0.77 |
| Labyrinth zone | 0.41 | 0.53 | 16.32 | **0.0002** | 0.01 | 0.94 |
| **Allopregnanolone** | | | | | | |
| Junctional zone | 1.12 | 0.30 | 0.77 | 0.39 | 0.50 | 0.48 |
| Labyrinth zone | 0.40 | 0.53 | 0.89 | 0.35 | 0.14 | 0.71 |
| **Estrone** | | | | | | |
| Junctional zone | 0.74 | 0.39 | 0.03 | 0.87 | 0.66 | 0.42 |
| Labyrinth zone | 0.33 | 0.57 | 0.20 | 0.66 | 0.42 | 0.52 |

Abbreviations: DOC: 11-deoxycorticosterone; DHC: 11-dehydrocorticosterone.

n = 12-15/diet/sex. Data were analyzed using 2-way ANOVAs. There was a significant main effect of Sex for androstenedione and testosterone in the labyrinth zone (bolded). Aldosterone and 17β-estradiol were below the lower limit of quantification.

**Supplementary Table 4: Additional steroids measured in the fetal blood and brain (ng/mL or ng/g).**

| | CON Male | | | HSD Male | | | CON Female | | | HSD Female | | |
|---|---|---|---|---|---|---|---|---|---|---|---|---|
| **Pregnenolone** | | | | | | | | | | | | |
| Blood | 8.08 | ± | 1.13 | 7.62 | ± | 0.78 | 8.49 | ± | 0.97 | 8.78 | ± | 0.94 |
| NAc | 35.55 | ± | 3.50 | 37.21 | ± | 2.96 | 37.97 | ± | 3.66 | 35.83 | ± | 2.44 |
| AMY | 36.99 | ± | 3.61 | 36.75 | ± | 3.09 | 43.13 | ± | 5.13 | 39.11 | ± | 3.47 |
| HYP | 36.12 | ± | 3.44 | 40.14 | ± | 2.97 | 38.96 | ± | 4.71 | 41.89 | ± | 5.01 |
| vHPC | 36.54 | ± | 4.39 | 38.23 | ± | 3.19 | 34.89 | ± | 3.32 | 43.42 | ± | 4.70 |
| VTA | 43.91 | ± | 3.94 | 47.25 | ± | 3.38 | 46.95 | ± | 6.65 | 52.39 | ± | 4.55 |
| **Progesterone** | | | | | | | | | | | | |
| Blood | 12.32 | ± | 1.13 | 12.83 | ± | 0.91 | 14.91 | ± | 0.99 | 12.63 | ± | 0.85 |
| NAc | 43.65 | ± | 3.72 | 54.81 | ± | 6.61 | 55.73 | ± | 4.74 | 49.84 | ± | 7.04 |
| AMY | 43.68 | ± | 4.18 | 43.84 | ± | 3.98 | 47.48 | ± | 3.07 | 41.43 | ± | 6.20 |
| HYP | 46.70 | ± | 4.66 | 50.73 | ± | 5.86 | 50.05 | ± | 4.17 | 49.28 | ± | 8.56 |
| vHPC | 39.99 | ± | 3.09 | 40.43 | ± | 3.23 | 40.47 | ± | 2.77 | 36.14 | ± | 3.53 |
| VTA | 41.74 | ± | 4.17 | 42.58 | ± | 4.24 | 48.85 | ± | 4.34 | 44.26 | ± | 6.43 |
| **DOC** | | | | | | | | | | | | |
| Blood | 4.70 | ± | 0.48 | 8.65 | ± | 2.20 | 6.13 | ± | 0.84 | 6.83 | ± | 0.79 |
| NAc | 2.73 | ± | 0.30 | 5.70 | ± | 1.53 | 3.52 | ± | 0.39 | 3.55 | ± | 0.45 |
| AMY | 3.00 | ± | 0.29 | 5.43 | ± | 1.51 | 3.33 | ± | 0.44 | 3.42 | ± | 0.34 |
| HYP | 2.93 | ± | 0.31 | 5.98 | ± | 1.66 | 3.44 | ± | 0.37 | 3.46 | ± | 0.46 |
| vHPC | 3.44 | ± | 0.40 | 5.82 | ± | 1.54 | 3.51 | ± | 0.43 | 3.78 | ± | 0.55 |
| VTA | 3.56 | ± | 0.36 | 6.06 | ± | 1.64 | 4.35 | ± | 0.54 | 4.28 | ± | 0.63 |
| **Corticosterone** | | | | | | | | | | | | |
| Blood | 218.12 | ± | 11.43 | 237.89 | ± | 19.46 | 252.37 | ± | 10.77 | 255.63 | ± | 17.32 |
| NAc | 30.78 | ± | 2.55 | 40.16 | ± | 5.28 | 41.73 | ± | 6.56 | 36.92 | ± | 4.18 |
| AMY | 26.85 | ± | 2.16 | 30.41 | ± | 3.97 | 31.27 | ± | 1.88 | 27.75 | ± | 2.11 |
| HYP | 19.61 | ± | 1.89 | 26.10 | ± | 4.61 | 27.80 | ± | 2.65 | 24.50 | ± | 2.56 |
| vHPC | 30.71 | ± | 2.33 | 34.37 | ± | 4.31 | 35.25 | ± | 3.95 | 31.34 | ± | 3.11 |
| VTA | 24.63 | ± | 2.59 | 27.69 | ± | 3.14 | 29.58 | ± | 2.69 | 25.46 | ± | 2.31 |
| **DHC** | | | | | | | | | | | | |
| Blood | 47.38 | ± | 3.55 | 46.77 | ± | 5.49 | 50.90 | ± | 4.49 | 45.24 | ± | 3.44 |
| NAc | 38.24 | ± | 3.15 | 38.83 | ± | 4.55 | 43.77 | ± | 3.48 | 34.48 | ± | 1.94 |
| AMY | 38.83 | ± | 2.24 | 40.06 | ± | 4.31 | 41.77 | ± | 2.58 | 35.93 | ± | 2.03 |
| HYP | 38.25 | ± | 3.01 | 41.44 | ± | 4.60 | 40.90 | ± | 3.22 | 35.48 | ± | 2.12 |
| vHPC | 36.56 | ± | 2.66 | 34.55 | ± | 2.95 | 36.46 | ± | 2.77 | 33.63 | ± | 2.99 |
| VTA | 37.31 | ± | 2.85 | 35.33 | ± | 3.05 | 42.55 | ± | 4.21 | 35.77 | ± | 2.73 |

| | CON Male | HSD Male | CON Female | HSD Female |
|---|---|---|---|---|
| **Androstenedione** | | | | |
| Blood | 1.18 ± 0.09 | 1.10 ± 0.07 | 0.63 ± 0.06 | 0.63 ± 0.07 |
| NAc | 2.12 ± 0.20 | 2.26 ± 0.25 | 1.28 ± 0.14 | 1.38 ± 0.20 |
| AMY | 2.02 ± 0.17 | 2.05 ± 0.20 | 1.22 ± 0.17 | 1.27 ± 0.15 |
| HYP | 2.05 ± 0.19 | 2.11 ± 0.23 | 1.12 ± 0.08 | 1.33 ± 0.17 |
| vHPC | 2.13 ± 0.16 | 1.97 ± 0.18 | 1.08 ± 0.11 | 1.10 ± 0.12 |
| VTA | 2.20 ± 0.16 | 2.24 ± 0.21 | 1.34 ± 0.15 | 1.36 ± 0.16 |
| **Testosterone** | | | | |
| Blood | 0.69 ± 0.06 | 0.67 ± 0.08 | 0.05 ± 0.00 | 0.07 ± 0.02 |
| NAc | 1.16 ± 0.10 | 1.43 ± 0.15 | 0.10 ± 0.02 | 0.13 ± 0.02 |
| AMY | 1.21 ± 0.09 | 1.25 ± 0.11 | 0.10 ± 0.03 | 0.14 ± 0.03 |
| HYP | 1.10 ± 0.11 | 1.34 ± 0.18 | 0.08 ± 0.02 | 0.11 ± 0.02 |
| vHPC | 1.19 ± 0.11 | 1.20 ± 0.14 | 0.09 ± 0.02 | 0.11 ± 0.02 |
| VTA | 1.19 ± 0.12 | 1.33 ± 0.16 | 0.10 ± 0.02 | 0.11 ± 0.02 |
| **Allopregnanolone** | | | | |
| Blood | 14.84 ± 1.09 | 13.82 ± 1.48 | 15.16 ± 1.11 | 14.05 ± 1.43 |
| NAc | 45.81 ± 2.47 | 41.44 ± 3.75 | 47.05 ± 3.51 | 42.97 ± 4.19 |
| AMY | 52.30 ± 3.77 | 39.50 ± 3.91 | 46.85 ± 3.74 | 42.60 ± 5.00 |
| HYP | 46.64 ± 3.31 | 39.89 ± 4.15 | 42.70 ± 3.60 | 38.31 ± 4.39 |
| vHPC | 48.51 ± 3.06 | 40.10 ± 4.25 | 45.10 ± 4.30 | 40.29 ± 5.23 |
| VTA | 59.04 ± 3.50 | 50.79 ± 5.50 | 55.49 ± 4.25 | 55.11 ± 5.82 |
| **Estrone** | | | | |
| Blood | 2.56 ± 0.14 | 2.62 ± 0.16 | 2.71 ± 0.20 | 2.68 ± 0.10 |
| NAc | 0.48 ± 0.04 | 0.43 ± 0.04 | 0.45 ± 0.04 | 0.40 ± 0.05 |
| AMY | 1.04 ± 0.11 | 0.88 ± 0.08 | 0.84 ± 0.10 | 0.78 ± 0.08 |
| HYP | 0.72 ± 0.04 | 0.77 ± 0.06 | 0.63 ± 0.03 | 0.70 ± 0.07 |
| vHPC | 0.48 ± 0.05 | 0.43 ± 0.04 | 0.44 ± 0.05 | 0.42 ± 0.04 |
| VTA | 0.42 ± 0.03 | 0.37 ± 0.05 | 0.28 ± 0.03 | 0.36 ± 0.03 |

Abbreviations: CON: control diet; HSD: high-sucrose diet; DOC: 11-deoxycorticosteorne; DHC: 11-dehydrocorticosterone; NAc: nucleus accumbens; AMY: amygdala; HYP: hypothalamus; vHPC: ventral hippocampus; VTA: ventral tegmental area.

n = 14-15/diet/sex. Data are presented as mean ± SEM. Data were analyzed using 2-way ANOVAs. See Supplementary Table 5 for statistical summary. Aldosterone and 17β-estradiol were below the lower limit of quantification.

**Supplementary Table 5: Statistical summary for all steroids in the fetal blood and brain.**

| | Diet | | Sex | | Diet × Sex | |
|---|---|---|---|---|---|---|
| | **F** | **p** | **F** | **p** | **F** | **p** |
| **Pregnenolone** | | | | | | |
| Blood | 0.00 | 0.95 | 0.89 | 0.35 | 0.01 | 0.92 |
| NAc | 0.01 | 0.94 | 0.03 | 0.87 | 0.36 | 0.55 |
| AMY | 0.30 | 0.59 | 1.19 | 0.28 | 0.23 | 0.63 |
| HYP | 0.84 | 0.36 | 0.11 | 0.74 | 0.13 | 0.72 |
| vHPC | 1.80 | 0.19 | 0.17 | 0.69 | 0.37 | 0.54 |
| VTA | 1.55 | 0.22 | 0.39 | 0.53 | 0.08 | 0.78 |
| **Progesterone** | | | | | | |
| Blood | 0.67 | 0.42 | 1.77 | 0.19 | 2.41 | 0.13 |
| NAc | 0.22 | 0.64 | 0.40 | 0.53 | 2.28 | 0.14 |
| AMY | 1.32 | 0.26 | 0.03 | 0.87 | 1.54 | 0.22 |
| HYP | 0.12 | 0.74 | 0.03 | 0.86 | 0.76 | 0.39 |
| vHPC | 0.38 | 0.54 | 0.36 | 0.55 | 0.57 | 0.45 |
| VTA | 0.15 | 0.70 | 0.82 | 0.37 | 0.31 | 0.58 |
| **DOC** | | | | | | |
| Blood | 2.40 | 0.13 | 0.76 | 0.39 | 0.38 | 0.54 |
| NAc | 2.39 | 0.13 | 0.01 | 0.92 | 2.52 | 0.12 |
| AMY | 1.49 | 0.23 | 0.11 | 0.75 | 0.59 | 0.45 |
| HYP | 1.45 | 0.23 | 0.28 | 0.60 | 2.07 | 0.16 |
| vHPC | 0.75 | 0.39 | 0.26 | 0.61 | 0.34 | 0.56 |
| VTA | 0.63 | 0.43 | 0.00 | 0.99 | 1.08 | 0.30 |
| **Corticosterone** | | | | | | |
| Blood | 0.29 | 0.59 | 4.10 | **0.05** | 0.29 | 0.59 |
| NAc | 0.22 | 0.64 | 0.86 | 0.36 | 1.84 | 0.18 |
| AMY | 0.17 | 0.68 | 0.68 | 0.41 | 1.29 | 0.26 |
| HYP | 0.12 | 0.73 | 2.41 | 0.13 | 2.73 | 0.10 |
| vHPC | 0.10 | 0.75 | 0.08 | 0.78 | 0.84 | 0.36 |
| VTA | 0.04 | 0.84 | 0.48 | 0.49 | 1.59 | 0.21 |
| **DHC** | | | | | | |
| Blood | 0.89 | 0.35 | 0.13 | 0.72 | 0.17 | 0.68 |
| NAc | 2.07 | 0.16 | 0.19 | 0.66 | 1.62 | 0.21 |
| AMY | 1.27 | 0.26 | 0.00 | 1.00 | 1.26 | 0.27 |
| HYP | 0.23 | 0.63 | 0.06 | 0.81 | 1.35 | 0.25 |
| vHPC | 0.73 | 0.40 | 0.03 | 0.86 | 0.02 | 0.89 |
| VTA | 1.69 | 0.20 | 0.79 | 0.38 | 0.45 | 0.51 |

| | Diet | | Sex | | Diet × Sex | |
|---|---|---|---|---|---|---|
| | **F** | **p** | **F** | **p** | **F** | **p** |
| **Androstenedione** | | | | | | |
| Blood | 0.29 | 0.59 | 50.09 | **<0.0001** | 0.39 | 0.53 |
| NAc | 0.35 | 0.56 | 18.42 | **<0.0001** | 0.01 | 0.91 |
| AMY | 0.01 | 0.91 | 22.21 | **<0.0001** | 0.04 | 0.85 |
| HYP | 0.64 | 0.43 | 23.70 | **<0.0001** | 0.19 | 0.67 |
| vHPC | 0.22 | 0.64 | 41.96 | **<0.0001** | 0.39 | 0.54 |
| VTA | 0.04 | 0.85 | 26.17 | **<0.0001** | 0.00 | 0.95 |
| **Testosterone** | | | | | | |
| Blood | 0.36 | 0.55 | 509.10 | **<0.0001** | 1.07 | 0.31 |
| NAc | 4.63 | **0.04** | 338.70 | **<0.0001** | 0.40 | 0.53 |
| AMY | 2.35 | 0.13 | 297.50 | **<0.0001** | 2.06 | 0.16 |
| HYP | 2.74 | 0.10 | 385.40 | **<0.0001** | 0.21 | 0.65 |
| vHPC | 0.91 | 0.34 | 407.10 | **<0.0001** | 1.12 | 0.29 |
| VTA | 0.00 | 0.95 | 249.10 | **<0.0001** | 0.20 | 0.66 |
| **Allopregnanolone** | | | | | | |
| Blood | 0.69 | 0.41 | 0.05 | 0.83 | 0.00 | 0.97 |
| NAc | 2.55 | 0.12 | 0.06 | 0.81 | 0.01 | 0.91 |
| AMY | 6.66 | **0.01** | 0.14 | 0.71 | 0.92 | 0.34 |
| HYP | 3.39 | 0.07 | 0.76 | 0.39 | 0.02 | 0.88 |
| vHPC | 2.43 | 0.13 | 0.14 | 0.71 | 0.18 | 0.67 |
| VTA | 2.01 | 0.16 | 0.01 | 0.92 | 0.89 | 0.35 |
| **Estrone** | | | | | | |
| Blood | 0.01 | 0.92 | 0.49 | 0.49 | 0.09 | 0.76 |
| NAc | 1.72 | 0.20 | 0.47 | 0.50 | 0.01 | 0.91 |
| AMY | 1.44 | 0.24 | 2.59 | 0.11 | 0.29 | 0.59 |
| HYP | 1.33 | 0.25 | 2.43 | 0.13 | 0.04 | 0.84 |
| vHPC | 0.82 | 0.37 | 0.46 | 0.50 | 0.02 | 0.88 |
| VTA | 0.22 | 0.64 | 4.68 | **0.04** | 6.98 | **0.01** |

Abbreviations: DOC: 11-deoxycorticosterone; DHC: 11-dehydrocorticosterone; NAc: nucleus accumbens; AMY: amygdala; HYP: hypothalamus; vHPC: ventral hippocampus; VTA: ventral tegmental area.

n = 14-15/diet/sex. Data were analyzed using 2-way ANOVAs. Bolded values indicate significant main effects of Diet or Sex, or a significant interaction ($p \leq 0.05$). Aldosterone and 17β-estradiol were below the lower limit of quantification.

**Supplementary Table 6: Statistical summary for DHC/corticosterone ratio in the fetal blood and brain.**

| | Diet | | Sex | | Diet × Sex | |
|---|---|---|---|---|---|---|
| | **F** | **p** | **F** | **P** | **F** | **p** |
| **DHC/corticosterone** | | | | | | |
| Blood | 1.55 | 0.22 | 1.56 | 0.22 | 0.01 | 0.92 |
| NAc | 2.68 | 0.11 | 0.42 | 0.52 | 0.19 | 0.66 |
| AMY | 0.29 | 0.59 | 1.20 | 0.28 | 0.05 | 0.82 |
| HYP | 0.73 | 0.40 | 4.40 | **0.04** | 0.90 | 0.35 |
| vHPC | 0.26 | 0.61 | 0.22 | 0.64 | 0.65 | 0.42 |
| VTA | 1.02 | 0.32 | 0.00 | 0.95 | 0.65 | 0.42 |

Abbreviations: DHC: 11-dehydrocorticosterone; NAc: nucleus accumbens; AMY: amygdala; HYP: hypothalamus; vHPC: ventral hippocampus; VTA: ventral tegmental area.

n = 14-15/diet/sex. Data were analyzed using 2-way ANOVAs. Bolded value indicates significant main effect. Aldosterone and 17β-estradiol were below the lower limit of quantification.

**Supplementary Table 7: Additional steroids measured in the amniotic fluid (ng/mL).**

| | CON Male | | | HSD Male | | | CON Female | | | HSD Female | | |
|---|---|---|---|---|---|---|---|---|---|---|---|---|
| Pregnenolone | 1.46 | ± | 0.29 | 1.95 | ± | 0.33 | 1.60 | ± | 0.23 | 1.70 | ± | 0.33 |
| Progesterone | 0.88 | ± | 0.21 | 0.91 | ± | 0.14 | 1.05 | ± | 0.16 | 0.86 | ± | 0.22 |
| Androstenedione | 0.16 | ± | 0.01 | 0.19 | ± | 0.02 | 0.11 | ± | 0.01 | 0.09 | ± | 0.01 |
| Testosterone | 0.05 | ± | 0.01 | 0.04 | ± | 0.01 | 0.01 | ± | 0.00 | 0.01 | ± | 0.00 |
| Allopregnanolone | 1.66 | ± | 0.29 | 1.75 | ± | 0.25 | 2.30 | ± | 0.44 | 1.96 | ± | 0.54 |
| Estrone | 0.65 | ± | 0.05 | 0.75 | ± | 0.07 | 0.73 | ± | 0.08 | 0.76 | ± | 0.15 |
| 17β-estradiol | 0.07 | ± | 0.01 | 0.09 | ± | 0.01 | 0.08 | ± | 0.01 | 0.09 | ± | 0.02 |

Abbreviations: CON: control diet, HSD: high-sucrose diet.

n = 14-15/diet/sex. Data were analyzed using 2-way ANOVAs. Data are presented as mean ± SEM. See Supplementary Table 8 for statistical summary.

**Supplementary Table 8: Statistical summary for all steroids measured in the amniotic fluid.**

| | Diet | | Sex | | Diet × Sex | |
|---|---|---|---|---|---|---|
| | **F** | **p** | **F** | **p** | **F** | **p** |
| Pregnenolone | 2.11 | 0.15 | 0.03 | 0.86 | 1.48 | 0.23 |
| Progesterone | 0.28 | 0.60 | 0.08 | 0.78 | 2.99 | 0.09 |
| DOC | 1.36 | 0.25 | 3.49 | 0.07 | 0.62 | 0.46 |
| Corticosterone | 0.00 | 0.95 | 0.04 | 0.84 | 2.10 | 0.15 |
| DHC | 4.08 | **0.05** | 3.81 | 0.06 | 2.82 | 0.10 |
| Androstenedione | <0.0001 | 1.00 | 27.98 | **<0.0001** | 0.84 | 0.36 |
| Testosterone | 0.72 | 0.40 | 159.60 | **<0.0001** | 0.42 | 0.52 |
| Allopregnanolone | 0.24 | 0.63 | 0.65 | 0.42 | 1.41 | 0.24 |
| Aldosterone | 0.13 | 0.72 | 0.02 | 0.89 | 0.34 | 0.56 |
| Estrone | 0.24 | 0.63 | 0.01 | 0.91 | 0.64 | 0.43 |
| 17β-estradiol | 1.96 | 0.17 | 0.04 | 0.84 | 0.73 | 0.40 |

Abbreviations: DOC: 11-deoxycorticosterone; DHC; 11-dehydrocorticosterone.

n = 14-15/diet/sex. Data were analyzed using 2-way ANOVAs. Bolded value indicates significant main effect.
